# The evolution of BDNF is defined by strict purifying selection and prodomain spatial coevolution, but what does it mean for human brain disease?

**DOI:** 10.1101/2022.01.21.477254

**Authors:** Alexander G Lucaci, Michael J Notaras, Sergei L Kosakovsky Pond, Dilek Colak

## Abstract

The mammalian gene Brain-Derived Neurotrophic Factor (BDNF) is an essential mediator of brain assembly, development, and maturation which has been implicated in a variety of brain disorders such as neurodevelopmental disorders (e.g. autism spectrum disorder), neuropsychiatric disorders (e.g. depression, PTSD, schizophrenia), and neurodegenerative disorders (e.g. Parkinson’s). Loss of BDNF during early development is embryonic lethal, and depletion of BDNF during adolescence or adulthood can result in disease-related neuropathology across a broad range of model organisms. In order to better understand the role of BDNF in disease, we seek to provide an evolutionary context to BDNF’s role within the brain by elucidating the molecular and genetic comparative history of BDNF across species. We conduct sequence alignment and phylogenetic reconstruction of the BDNF gene across a diverse selection of over 160 mammalian species spanning ∼177 million years of evolution. Selective evolutionary change was examined via several independent computational models of codon evolution including FEL, MEME, and BGM. We report strict purifying selection in the main functional domain of BDNF (NGF domain, essentially comprising the mature BDNF protein). Specifically, we discover 6 sites in our homologous alignment which are under episodic selection in the early regulatory region of BDNF (i.e. the prodomain) and 23 pairs of coevolving sites that are a part of complex spatial relationships that are distributed across the entire BDNF gene. Thus, we propose that our discovery of both local and distal sites of co-evolution within the pro- and mature-domains of BDNF that likely reflect the evolutionary fine-tuning of BDNF’s unique and complex regulatory capacities whilst also retaining it’s core yet diverse ontogenic functionality within the central nervous system. This discovery consequently supports the idea that the BDNF prodomain is more prone to change than the mature domain, however the fact that this region has also been subject to negative purifying selection also highlights genetic sensitivity and thus partially explains the prodomain’s disease relevance (e.g. Val66Met and other variants) to numerous neuropsychiatric disorders.

**HIGHLIGHTS:**

- We extracted coding sequences for Brain-Derived Neurotrophic Factor (BDNF) from over 160 mammalian genomes that span approximately ∼177 million years of evolution.
- We observe strict purifying selection in the main functional domain (NGF) of the BDNF gene in mammals.
- We observe novel results with 6 sites in our homologous alignment which are under episodic selection in the early regulatory region of BDNF (i.e. the prodomain).
- We observe 23 pairs of coevolving sites within BDNF. Many of which are a part of complex spatial relationships and are distributed across the entire BDNF gene.
- These data define exactly how “BDNF is highly conserved” by defining exactly where and how the mammalian BDNF has evolved, confirming the widespread belief that the BDNF prodomain is more prone to change than the mature BDNF protein.

## INTRODUCTION

Brain-Derived Neurotrophic Factor (BDNF) is an ubiquitously studied molecule in modern neuroscience [1]. BDNF is a neurotrophin that binds with high affinity to its cognate tyrosine kinase receptor, TrkB [2], to elicit rapid induction of synaptic plasticity [3–5] and neuronal spine remodeling [6, 7]. Additionally, BDNF has been implicated in a variety of brain disorders [1], including depression [8–10], PTSD [11–14], schizophrenia [9, 15–17], Parkinson’s disease [18, 19], and autism spectrum disorders [20–22] amongst many more. BDNF has correspondingly been the primary target, or an ancillary factor, of many novel therapeutics including small molecule mimetics [23, 24] and existing drugs (e.g. antidepressants [25, 26]). Yet, nascent research has provided the humbling reminder that much remains to be discovered about BDNF. In recent years, new BDNF ligands have been discovered [27], new receptor interactions unveiled [27, 28], and mechanisms of behavioral function unlocked [7]. This is a timely reminder that while BDNF has remained a seminal molecule of interest across the broader neuroscience literature, much remains to be discovered about its origins, evolution, function, and disease relevance.

### A Primer of the Molecular Biology of BDNF and its Functional Topology

BDNF is encoded by the *BDNF* gene [29], whose expression is regulated in humans by an antisense-gene (*BDNF-AS*) that can form RNA-duplexes to attenuate translation [30]. Thus, the natural antisense for BDNF is capable of directly downregulating endogenous expression on demand [31]. The *BDNF* gene in humans comprises 11 exons [30] and can produce at least 17 detectable transcript isoforms [29]. Different transcripts are induced in response to activity and/or cellular states, allowing the *BDNF* gene to adjust to environmental stimuli and potential selection pressures. However, all transcripts ultimately yield a singular preproBDNF protein that (prior to intracellular processing, cleavage, and transport) can be partitioned into three domains [11, 29]: a signal peptide, a prodomain, and the mature domain. The signal domain is only 18 amino acid residues long, possessing ambiguously defined functionality, and the majority of BDNFs functional outputs reflect sequence specificity to the prodomain and mature domain. The BDNF prodomain encodes binding sites for intracellular transport of both *BDNF* mRNA [32] and BDNF protein [33], and contains numerous post-translational modification sites [29]. The BDNF prodomain is also the resident location of a widely studied Single Nucleotide Polymorphism (SNP) in neuroscience (Val66Met, or rs6265) [1], and the Furin consensus sequence (Arg 125) for cleavage to its mature form (including by plasmin [34]). The prodomain is composed of 110 amino acids within the N-terminus, and must be processed via proteases to generate mature BDNF [5]. The mature domain of BDNF is composed of, almost exclusively, the Nerve growth factor (NGF) domain and is responsible for the canonical trophic actions associated with BDNF, e.g., long-term potentiation, rapid acting antidepressant effects. Following intracellular handling, processing, and transport, the preproBDNF isoform is cleaved to yield the mature BDNF peptide (which only contains the mature NGF domain). For many years the prodomain was thought to be cleaved following transport and thus destined for degradation. However, recent work has shown that the cleaved prodomain can be secreted and bind as a ligand to novel receptors (e.g., SorCS2) [27]. Thus, the BDNF prodomain can accordingly influence brain circuits as well as behavior [7]. For a comprehensive, detailed, analysis of the various intricacies of the BDNF gene, protein, and its regulation, more information is provided in [29].

### The Conservation of BDNF & Neurotrophins

One of the interesting curiosities surrounding BDNF is its relationship to other neurotrophic (NT) growth factors, comprising NGF, NT-3, NT-4. Thus, neurotrophins retain some intercalated functionality. Each share some commonalities in structure (pre-, pro-, and mature-domains) [29], post-translational modification potential (e.g. glycosylation [35]), as well as catalytic processing, trafficking, and composition [36]. Specifically, neurotrophins share approximately 50% sequence homology [29], and a comparison of domains and motifs reveals that each comprises a prototypic NGF-domain as the principal component of the mature pro-growth peptide for each factor (see PFAM database [37]). While each neurotrophin elicits functionality via binding to cognate receptors, neurotrophins also exhibit cross-affinity amongst neurotrophin receptors [38] presumably due to their high rates of structural homology. Not surprisingly then, there is some redundancy in the pro-trophic effects of neurotrophins, yet each still maintains nuanced functionality which remains specific to each factor during central nervous system development [39]. Differences in the evolution and temporal dynamics of regulatory sequences, which target gene-products to specific destinations within cell-compartments (e.g. dendrites) [40] or to processing routes (e.g. the activity-dependent release pathway) which alter secretory dynamics and/or bioavailability [41], likely contribute to both similarities and differences between neurotrophins. However, almost nothing is known of how the BDNF prodomain has evolutionarily adapted to specifically regulate BDNF dynamics. While evolution has almost certainly shaped the sequences, structure, and function of BDNF, the modeling of such remains relatively unexplored but could provide important insight into the phylogenetic evolutionary history of BDNF, it’s selection pressure sensitivity across lineages, and quantitative metrics of evolutionary change across species.

### Purpose of this Study

Here, we use computational methods to explore the comparative evolutionary genomics of the neurotrophic factor BDNF. By reconstructing phylogenetic trees of BDNF in mammals (*Mammalia*), we utilize sequence alignments of over 160 species to determine unique genomic attributes of BDNF. In specific we investigate which sites in BDNF are subject to pervasive (i.e., consistently across the entire phylogeny) diversifying selection (FEL) or pervasive/episodic (i.e., only on a single lineage or subset of lineages, diversifying selection (MEME). Likewise, utilizing multiple models for the inference of selective pressure and the evaluation of evolutionary change, we identify novel sites within the BDNF prodomain and mature peptide coding regions that are susceptible to synonymous and nonsynonymous changes. Additionally we investigate which sites in BDNF may be coevolving (BGM). Taken together, these computational evolutionary analyses provide an important context as to the origins and sensitivity of genetic changes within the BDNF gene, which may be important for providing insight into genetic risk factors linked to disease in humans.

## RESULTS

We find that unique evolutionary pressures have shaped the BDNF gene across time. Mostly, these forces have operated through strict purifying selection. Of note, BDNF elicits tight regulation and specific functionality that can be separated from other neurotrophins, yet these growth factors remain closely related in their structure and sequence, especially in the conserved NGF domain.

### Evolutionary History of Mammalian BDNF

Prior to conducting our primary evolutionary analysis, we ported our mammalian species into a platform (*timetree.org*, see [42, 43]) to examine the epoch events that may have influenced the analysis described here. This was an important pre-analysis step to frame the age of our genomes, and the broad-strokes evolutionary pressures that these species have been exposed to (which, in theory, could contribute to subsequent purifying selection and coevolution analyses). As expected, this revealed BDNF as an ancient gene that has been preserved throughout the mammalian lineage and has both survived and been shaped under all major evolutionary events of the past ∼177 million years (data not shown). For species within our data-set we identified several examples of species-level evolutionary epochs that cross-referenced with major earth events (e.g. bottleneck events) that have historically been believed to drive evolutionary adaptation. This included major geologic periods that are cross-referenced against earth impacts, oxygenation changes across time, atmospheric carbon dioxide concentrations, and solar luminosity. This indicates that even under extreme evolutionary pressures, the BDNF gene has exhibited (relatively speaking) very specific adaptation events (see results below) over millions of years in mammals. This tracks with the idea that “old genes” tend to be highly conserved, evolve more slowly, and therefore are more likely to exhibit both specific and selective changes as opposed to more dramatic permutations (e.g. gene duplications etc.).

### Predominant Purifying Selection in BDNF

A common approach to gain an increased understanding of the evolutionary forces that have shaped proteins is to measure the omega ratio ω consisting of the non-synonymous (β or dN) and synonymous (α or dS) substitution rates with ω = β/α for each site in a particular gene of interest [47]. We define two major substitional changes for the amino acid being coded for at each site: synonymous changes, which keep the same amino acid coded for at a particular site and nonsynonymous changes, which change the amino acid coded for at that particular site. Non-synonymous changes can have strong influences on the structural, functional, and fitness measures of an organism. This is in contrast to synonymous changes which leave the amino acid at a particular site unchanged, but can confer weak fitness effects through the emergent properties of codon usage bias, mRNA structural stability, translation and tRNA availability. However, synonymous changes are typically understood to represent neutral selection acting on coding sequences, and provides a baseline rate against which non-synonymous evolutionary rates can be compared. The omega ratio ω of relative rates of non-synonymous and synonymous substitutions is a common measure in evolutionary biology of the selective pressure acting on protein coding sequences. These estimates provide increased information availability as to the type of selection (positive, with omega > 1 or negative, with omega < 1, or neutral with omega = 1) that has acted upon any given set of protein-coding sequences.

As FEL analysis is a sensitive measure of *negative* (purifying) selection, for our FEL analysis, we observe a predominant amount of purifying selection (over 66% of sites, 174 sites out of 261; Table S1) in our recombination free alignment for BDNF. The dN/dS estimates for the entire alignment were plotted including 95% lower- and upper-bound estimates (see Figure 1 or Table S1). Overwhelmingly, the mature NGF-domain of the BDNF exhibited evidence of greater pervasive negative purifying selection relative to the prodomain region of BDNF. Thus over the evolutionary history in mammalia, negative selection has predominantly occurred in the regions of BDNF that encode the functional mature protein that binds TrkB to elicit neurotrophic effects. The mature domain of BDNF has exhibited remarkable conservation across innumerous epochs defined by rapid evolutionary adaptation (see Figure 1) in other genes and species.

**Figure 1.**
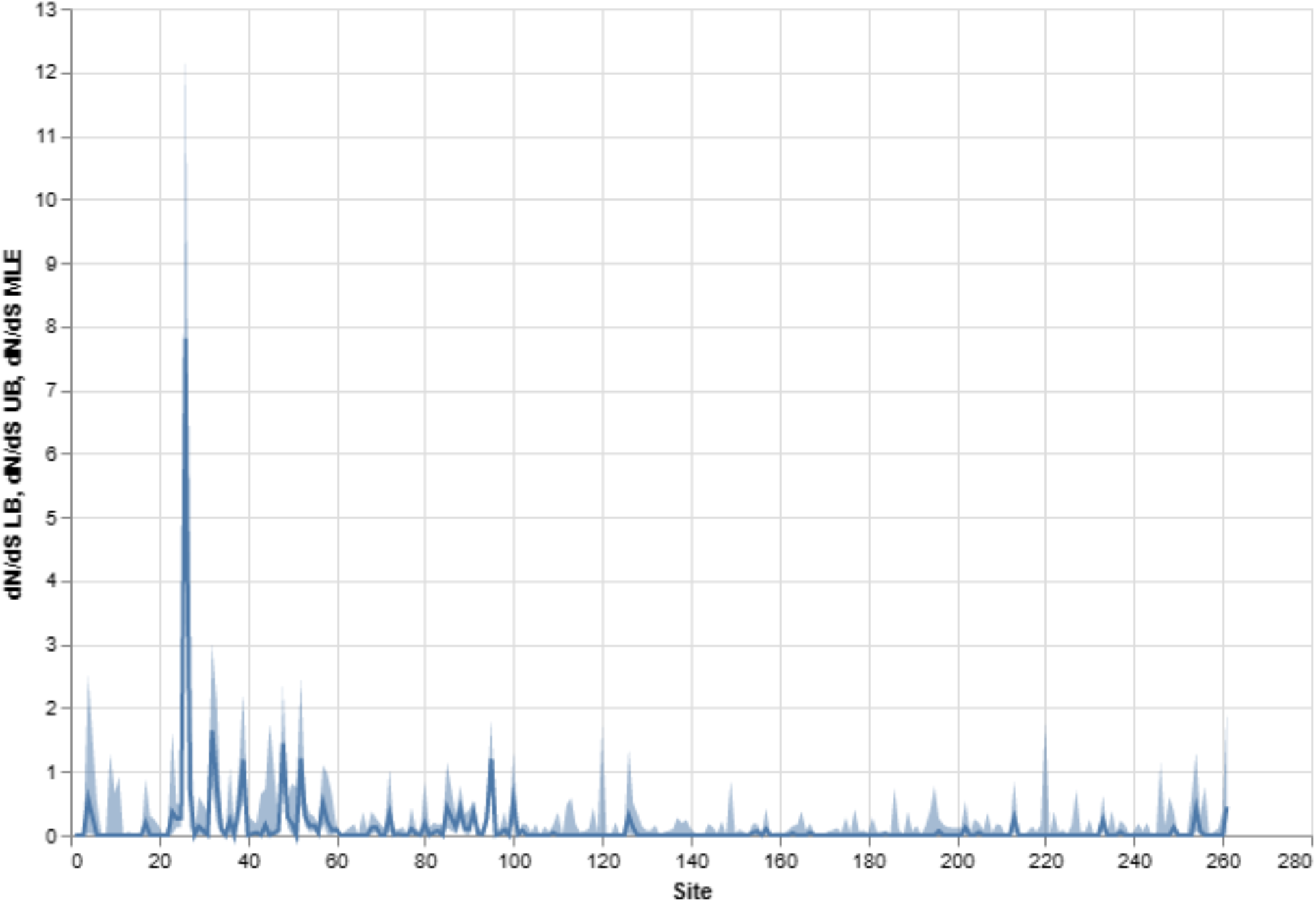
The FEL analysis of the BDNF gene found 174 of 261 (66.7%) sites to be statistically significant (LRT p-value <= 0.1) for pervasive negative (purifying) selection. We plot the estimated values of omega (dN/dS) for each site in the alignment. Additionally, we plot 95% confidence intervals (CI) for each site. These results are also available in Table S1. We observe a high degree of strict purifying selection in the Human NGF region. The region for Human NGF corresponds to alignment sites 144-254 (<u>NP_001700.2</u>, and https://github.com/aglucaci/AnalysisOfOrthologousCollections/blob/main/tables/BDNF/BDNF_AlignmentMap.csv). This alignment of BDNF across all selected species (Mammalia, see Table 1) reveals a site-specific positive/adaptive diversifying selection and negative purifying selection. The thick line represents the point estimate (i.e. the evolutionary pressure) and the shadings reflect 95% confidence intervals which relate to the upper and lower bound of the point estimates. As shown, the prodomain sites exhibit more pervasive/episodic and positive/diversifying evolutionary selection, consistent with the fact that more disease associated SNPs occur in this topological region of the BDNF gene in humans (i.e. early prodomain mapping not further shown due to nuanced variation across mammalian species).

**Table 1.**
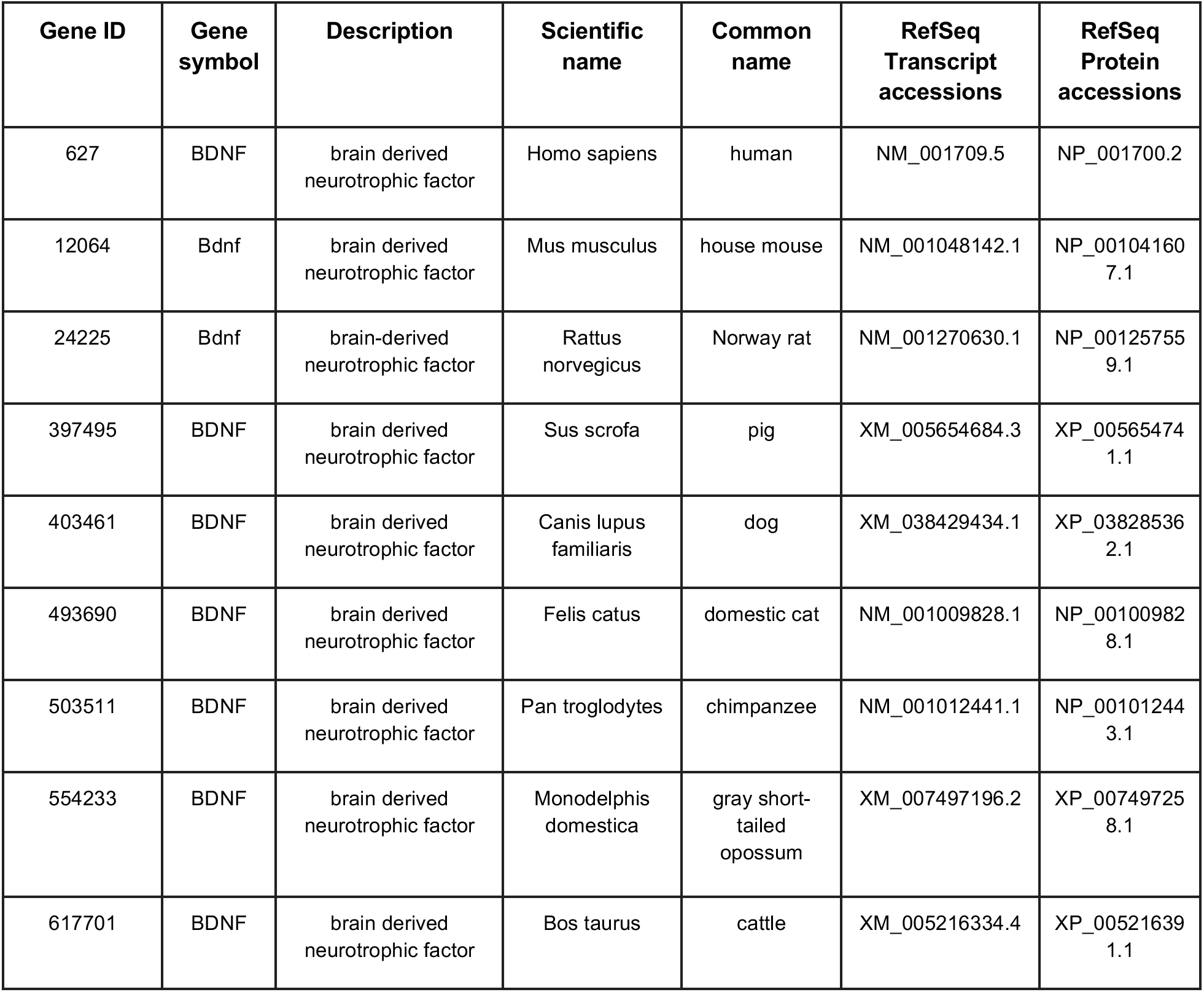
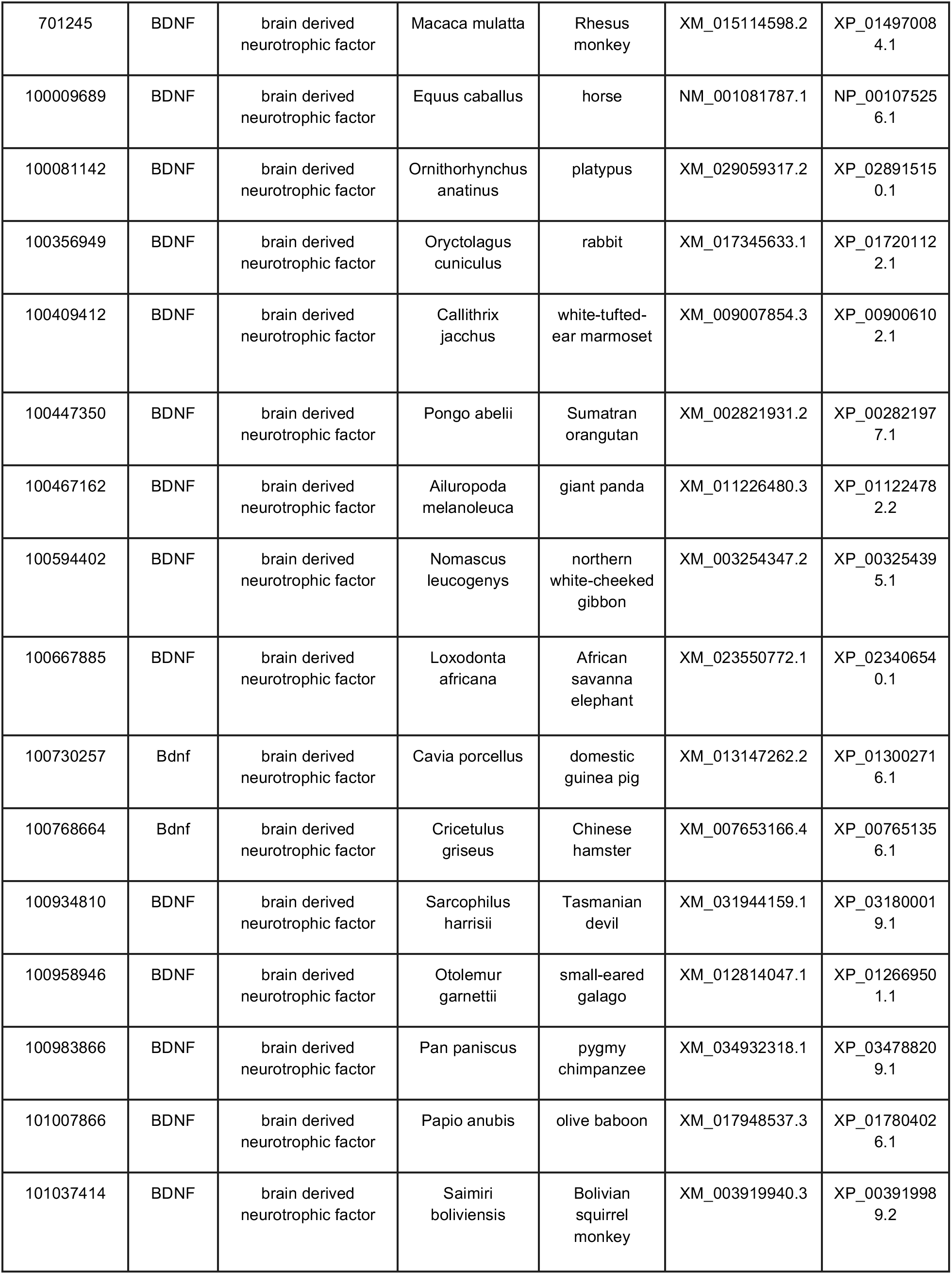
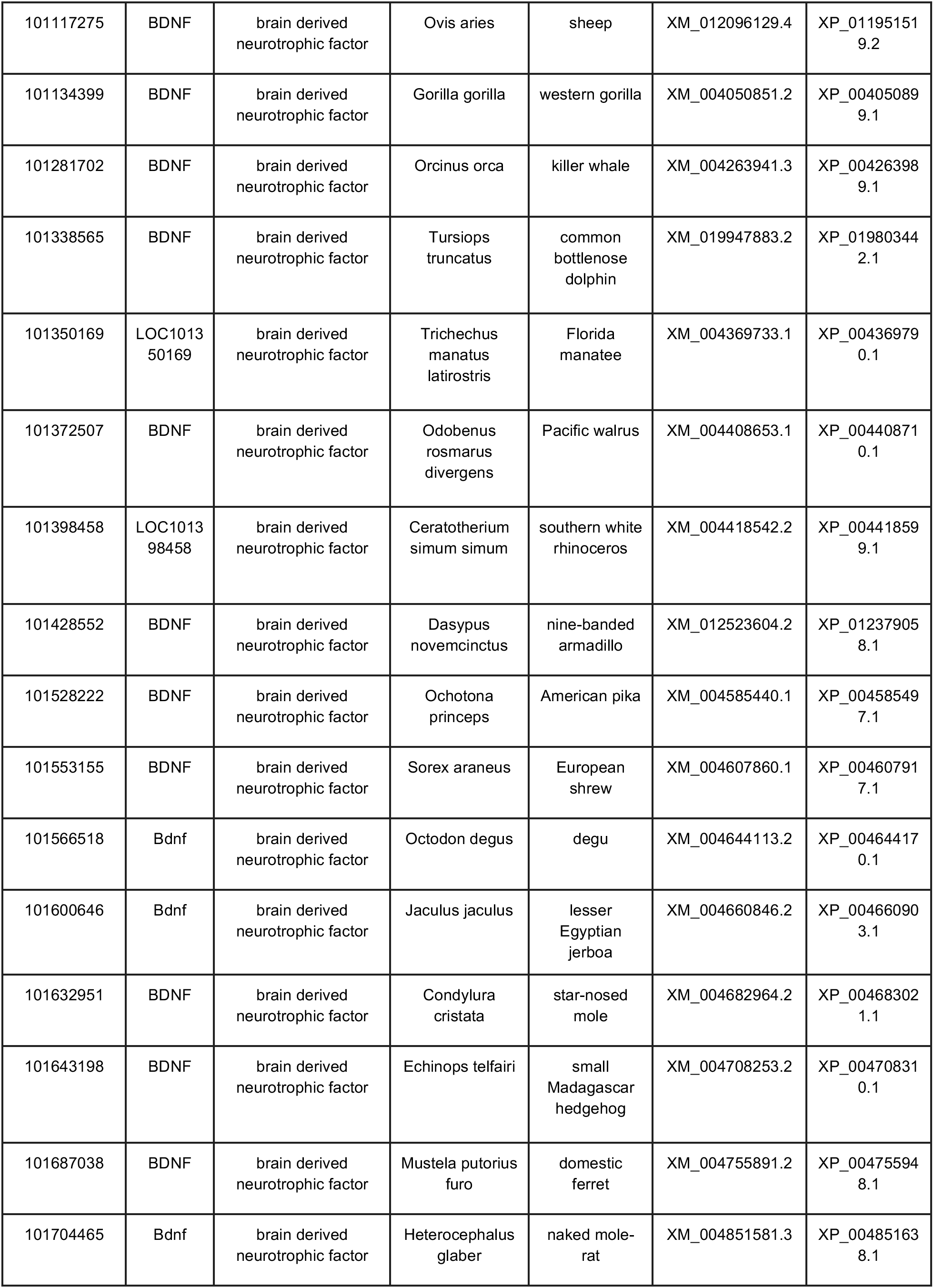
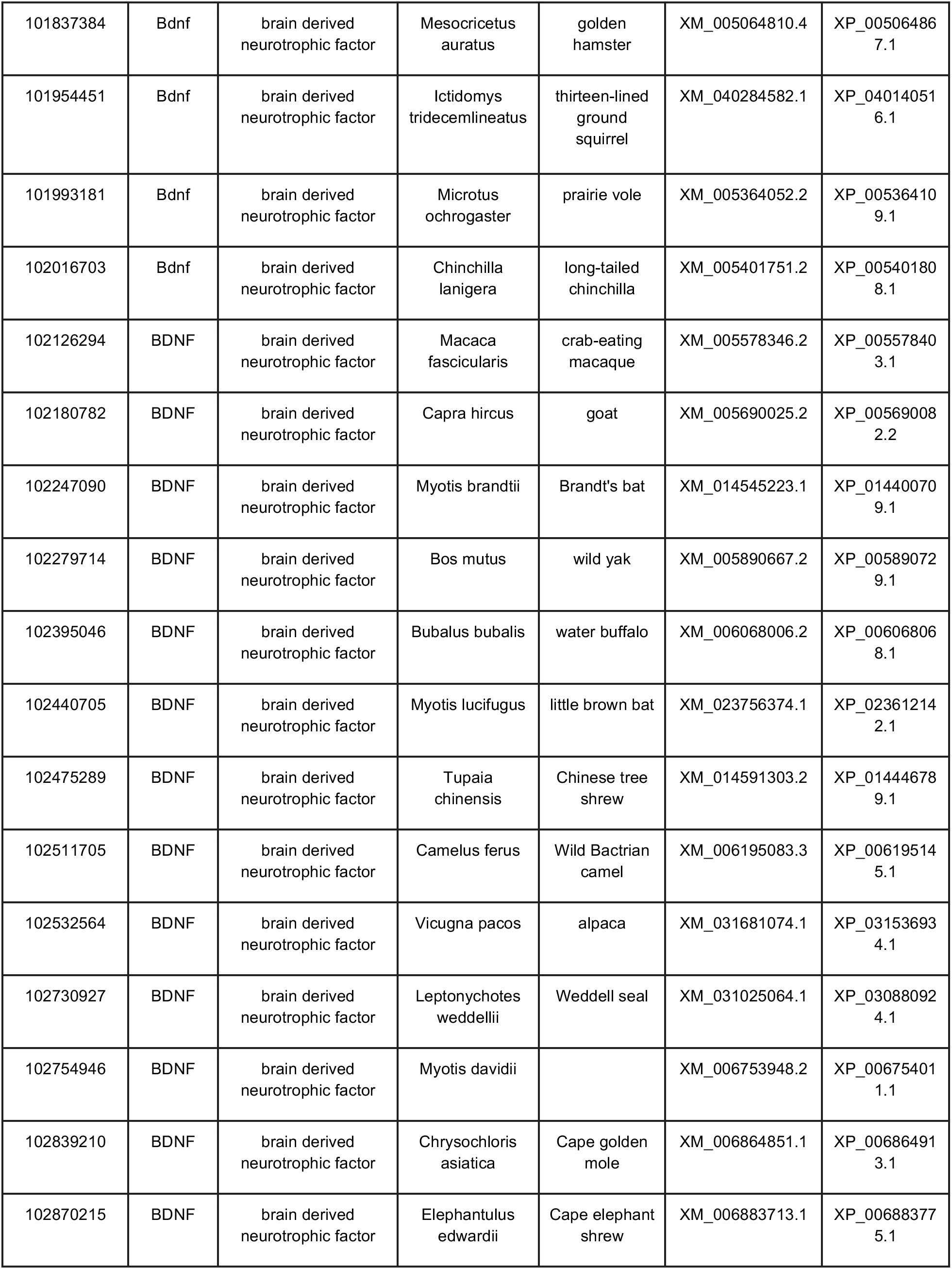
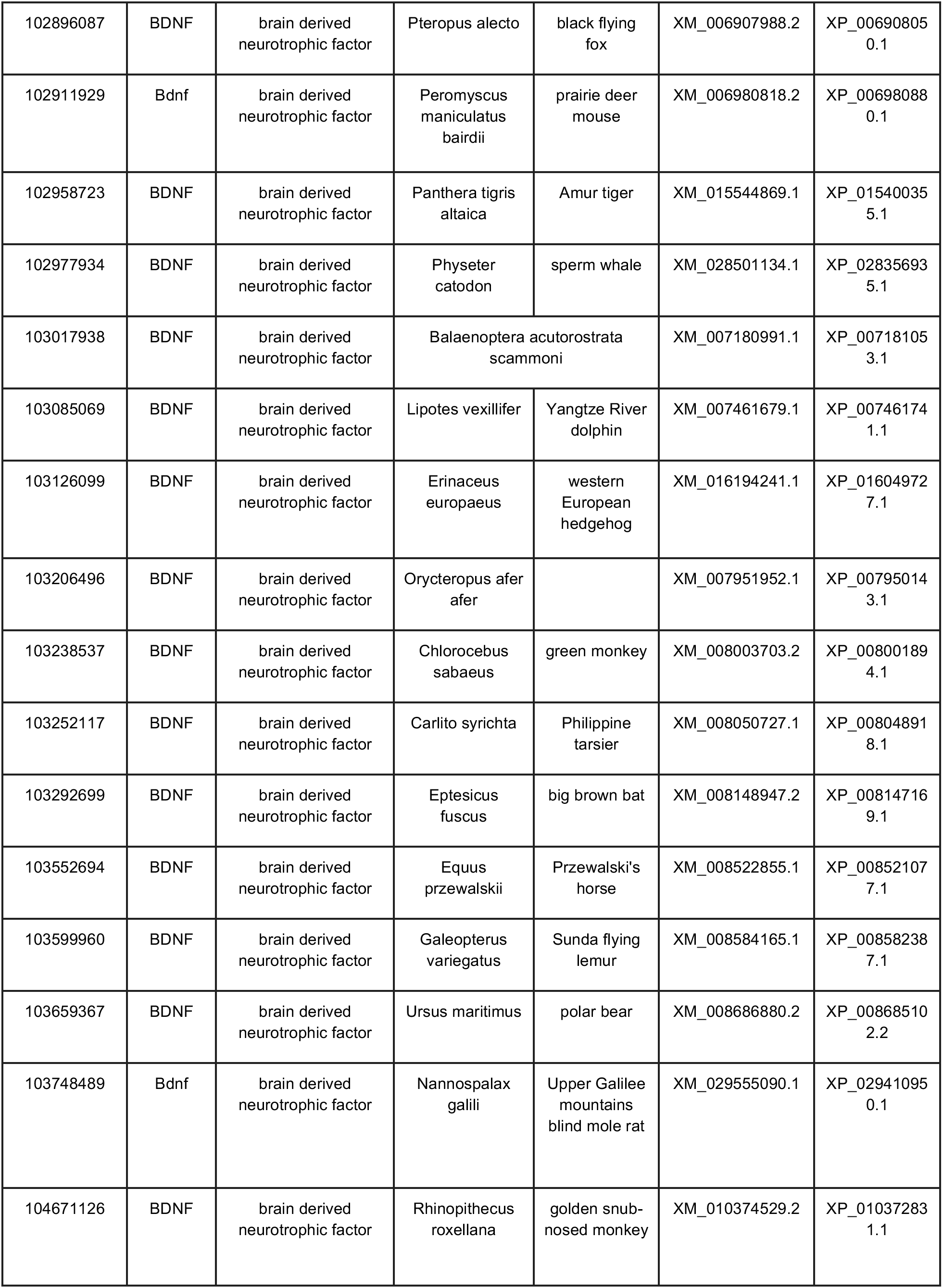
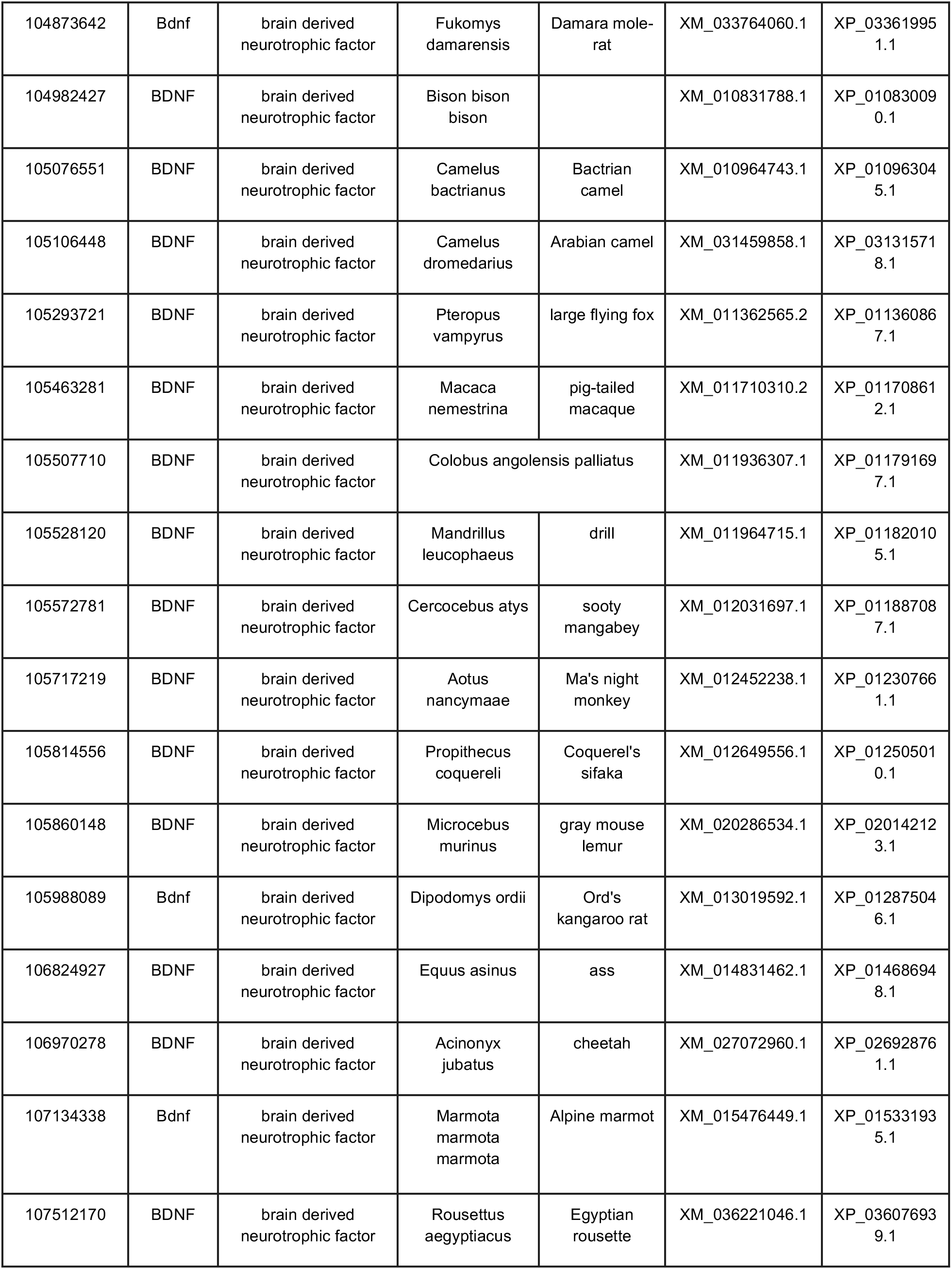
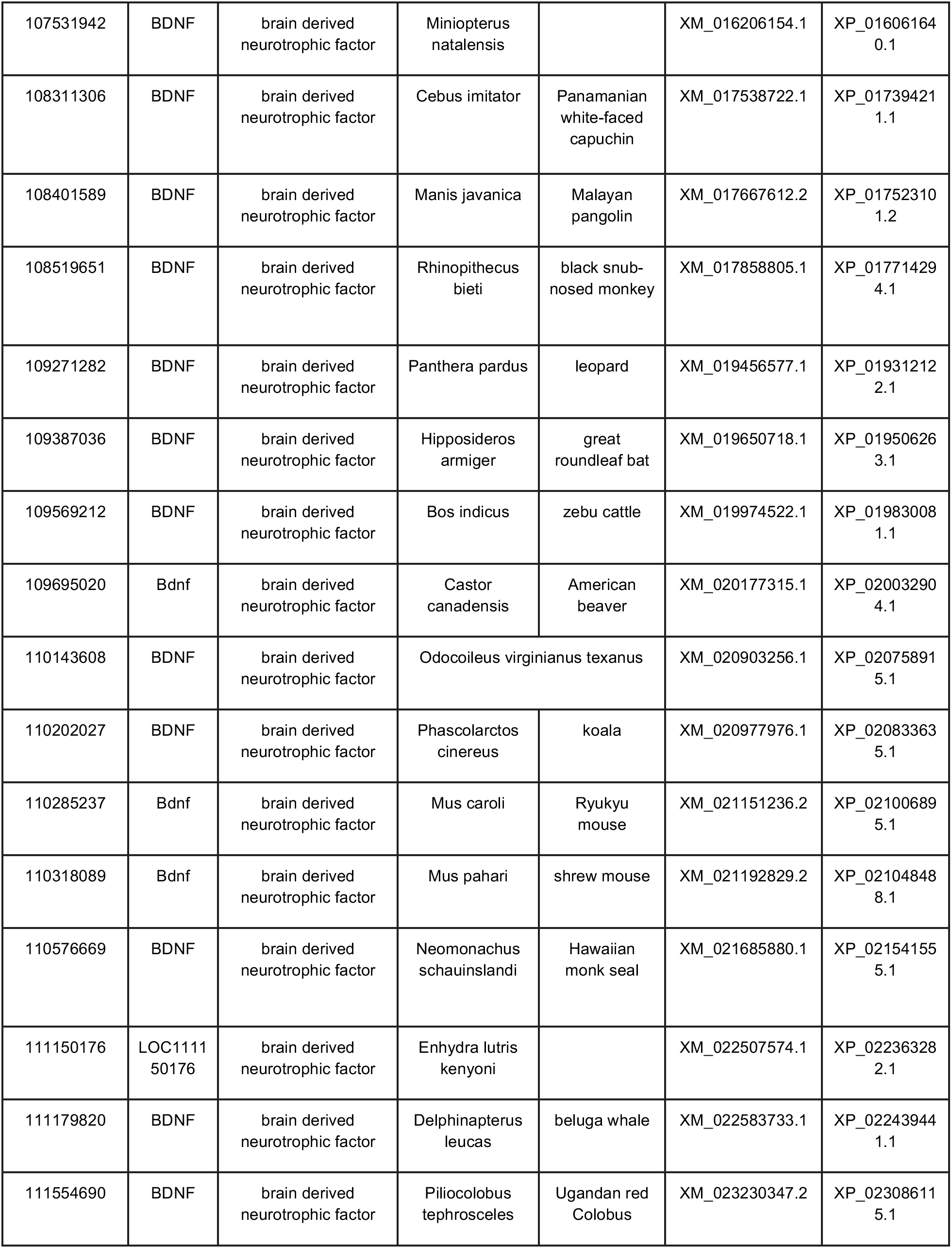
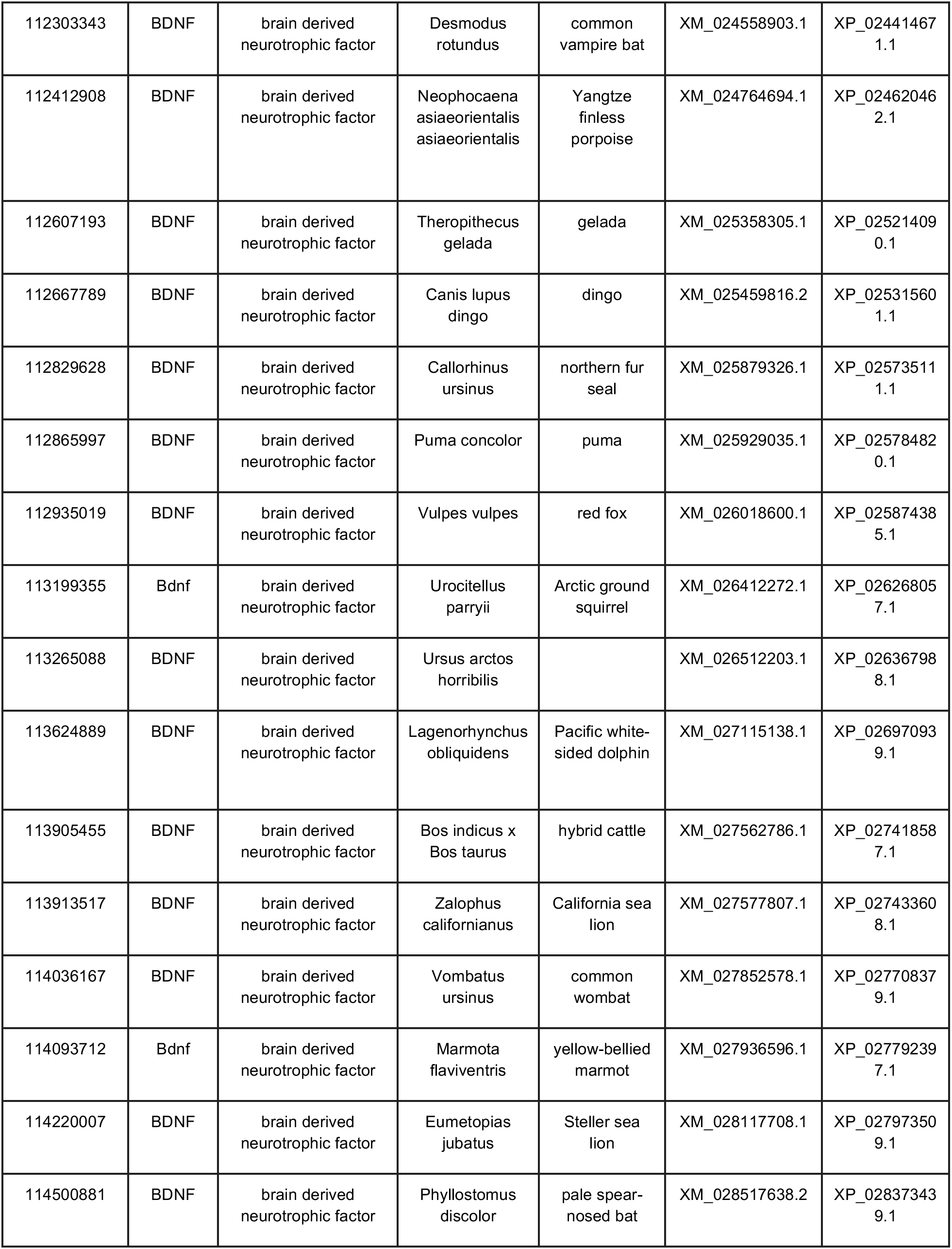
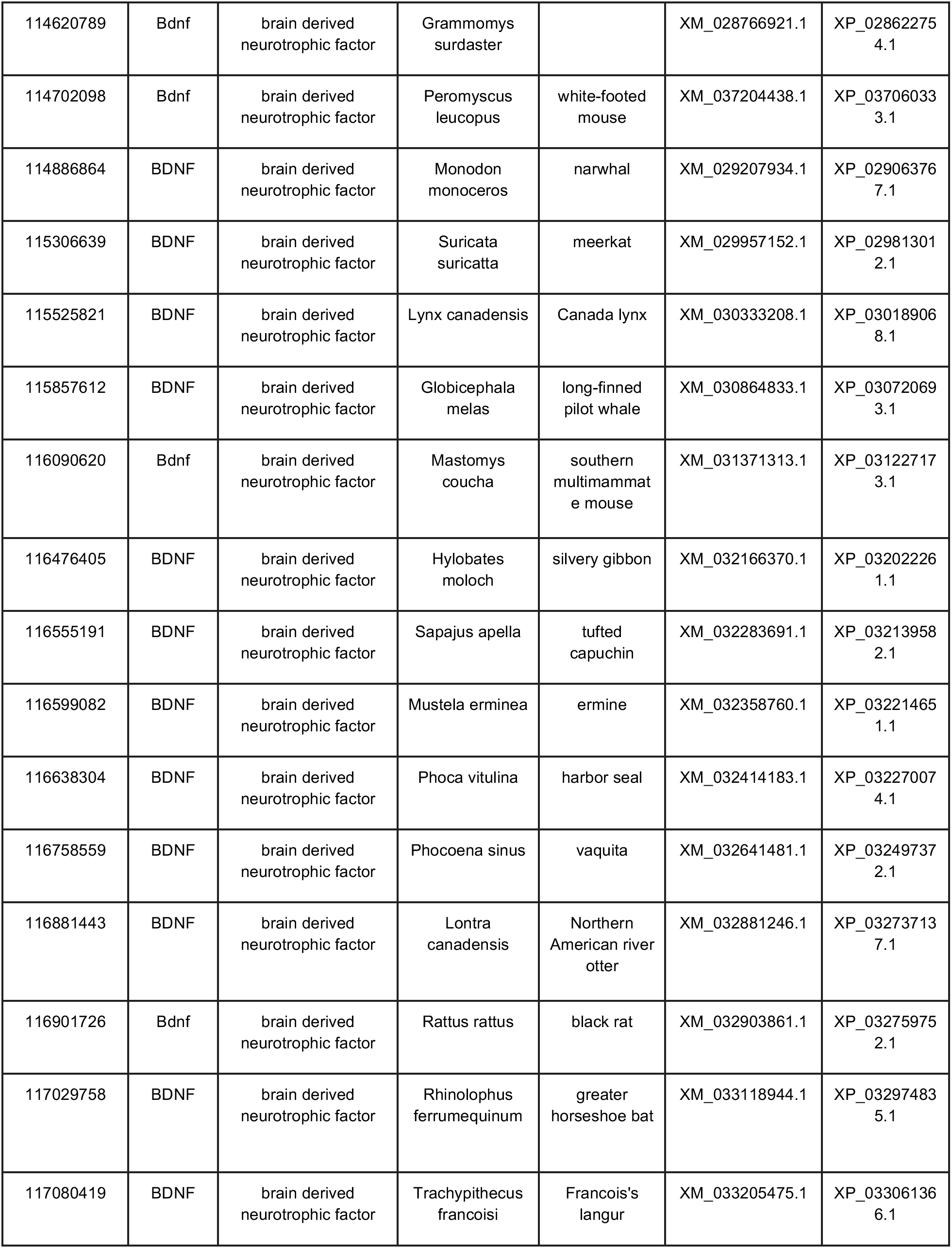
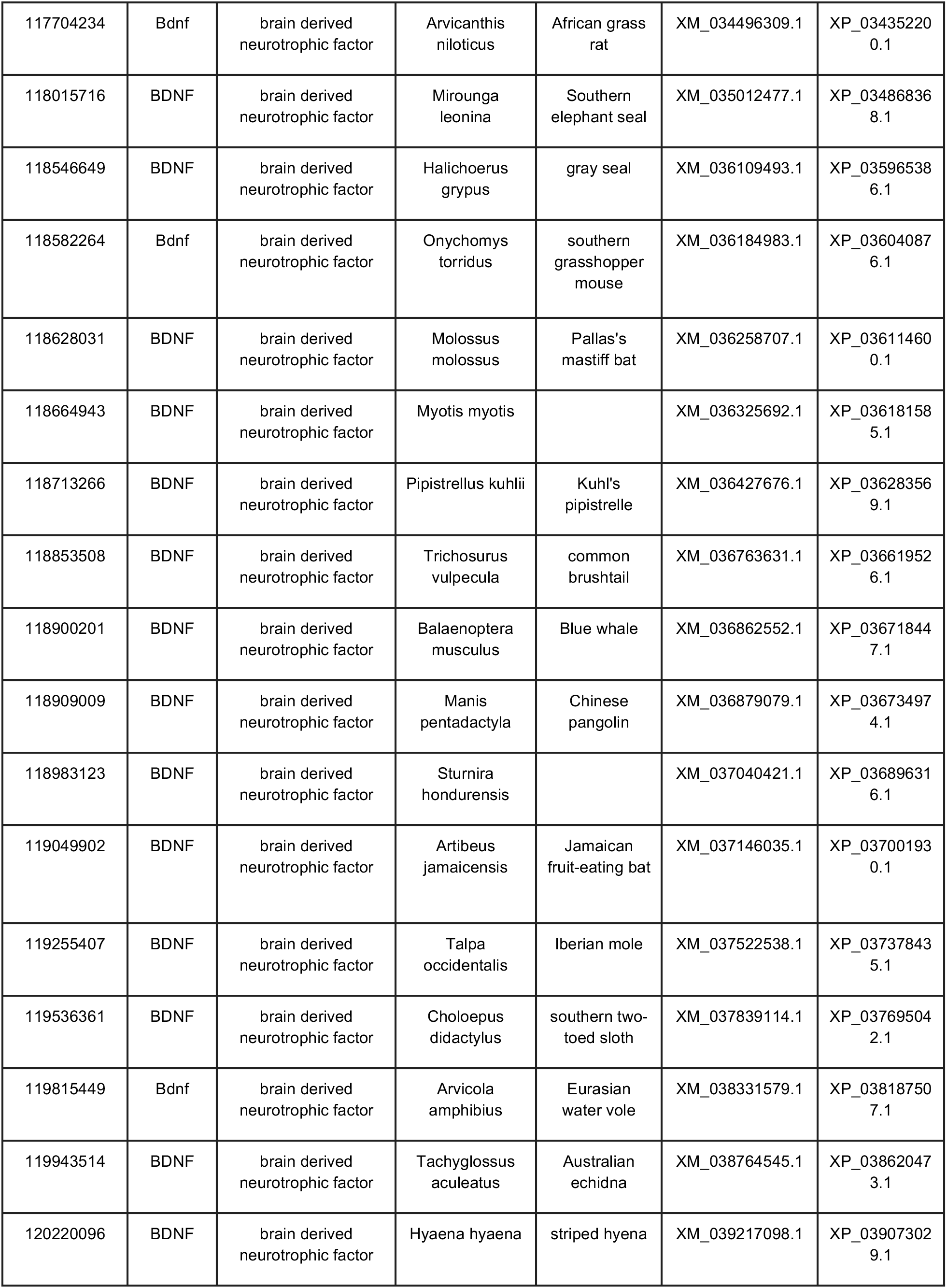
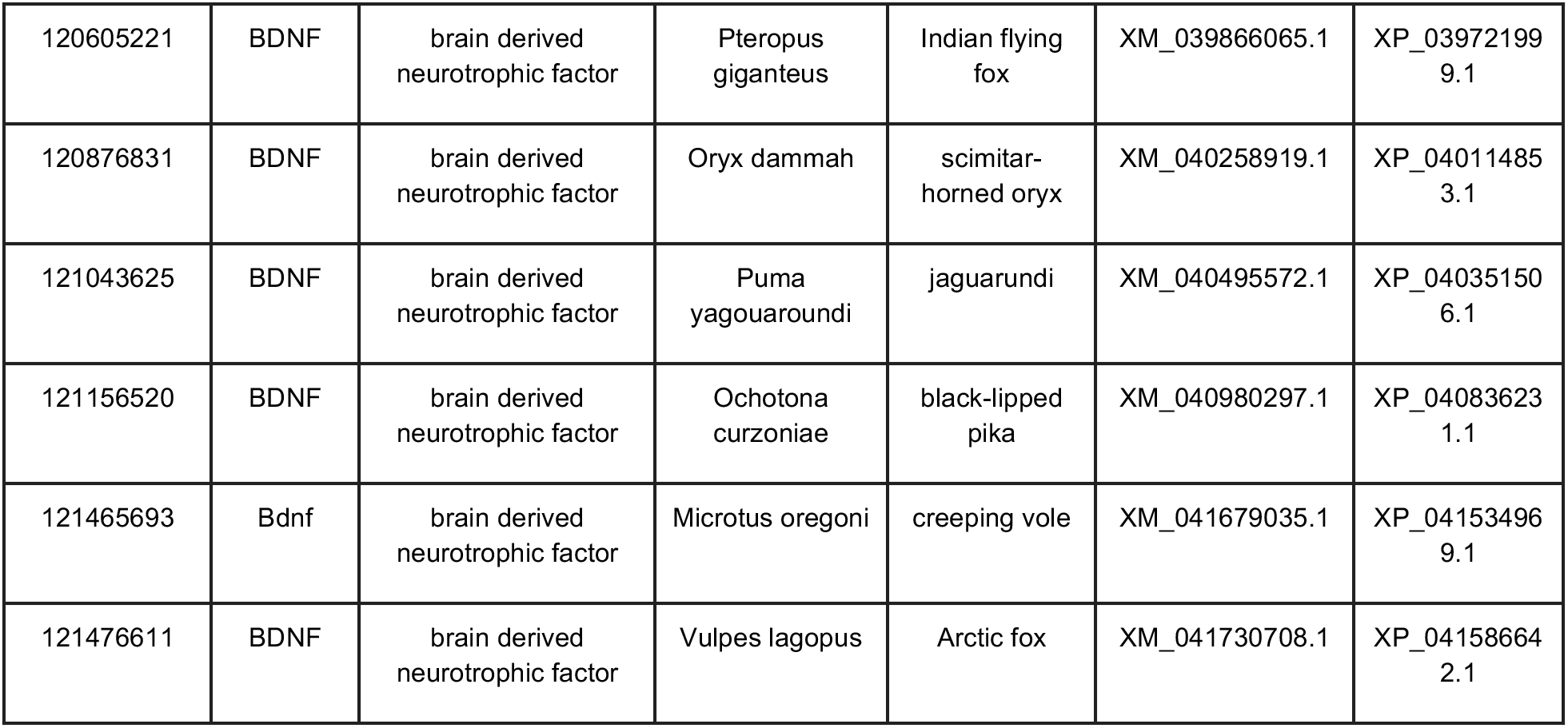
Tabulation of Species included in our analysis, comprising NCBI ortholog gene IDs, symbols, mammalian species, common name, and RefSeq accessions.

### Specific Sites that are Evolving Non-Neutrally

To examine specific sites for episodic adaptive evolutionary selection, we utilized an algorithm known as MEME which is fundamentally similar to our FEL analysis (described above) except that it applies a more sensitive method for the detection of both pervasive (persistent) and episodic selection (transient selection occurring only on one or a subset of branches in the phylogenetic tree) as compared to only pervasive selection which occurs across all branches of the phylogenetic tree. Essentially, only a subset of the lineages (i.e. species) are affected allowing for a more granular/sensitive method of detecting selection (whereas FEL is better geared towards *broad* changes). This analysis revealed that for all sites, only 2.3% (6 of 261; see Table 2) exhibit evidence for episodic diversifying selection (i.e. positive selection) in at least one branch within the phylogeny. Spatially, these mutations occur outside of the NGF functional region of BDNF. Further, this result is essentially relevant as the MEME analysis is a sensitive measure of episodic selection. The sites we observe as statistically significant are as follows: 26, 27, 30, 38, 249, 254. For comparison, these specific sites were re-aligned to the respective human sites with indel (insertion/deletion) events accounting for any respective discrepancy in specific site numbers. When mapping these sites to the human BDNF coordinate system, they correspond to: 26, 27, 29, 36, 238, 240, respectively.

**Table 2.** MEME analysis of the BDNF gene found 6 of 261 (2.3%) of sites to be statistically significant (LRT p-value <= 0.1).

| # | <u>Codon Site</u> | <u>Human Codon Site</u> | <u>alpha</u> | <u>beta-</u> | <u>p-</u> | <u>beta+</u> | <u>p+</u> | <u>LRT</u> | <u>p-value</u> | <u># branches under selection</u> | <u>MEME LogL</u> | <u>FEL LogL</u> | <u>Omega</u> |
| --- | --- | --- | --- | --- | --- | --- | --- | --- | --- | --- | --- | --- | --- |
| 1 | 26 | 26 | 0.14 | 0.00 | 0.02 | 1.12 | 0.98 | 6.89 | 0.01 | 9 | -86.53 | -86.53 | 7.9 |
| 2 | 27 | 27 | 0.25 | 0.08 | 0.99 | 28.91 | 0.01 | 4.30 | 0.05 | 1 | -39.25 | -37.01 | 117.6 |
| 3 | 30 | 29 | 0.00 | 0.00 | 0.00 | 0.38 | 1.00 | 4.63 | 0.05 | 6 | -35.88 | -35.88 | inf |
| 4 | 38 | 36 | 0.72 | 0.00 | 0.93 | 8.07 | 0.07 | 3.80 | 0.07 | 5 | -69.22 | -66.28 | 11.3 |
| 5 | 249 | 238 | 0.92 | 0.00 | 0.99 | 203.11 | 0.01 | 8.25 | 0.01 | 1 | -50.74 | -44.13 | 221.6 |
| 6 | 254 | 240 | 0.23 | 0.00 | 0.99 | 1357.12 | 0.01 | 21.81 | 0.00 | 1 | -44.50 | -31.95 | 5850.4 |

### Evidence of Coevolutionary Forces

To examine the coevolution of sites, i.e. if one particular amino acid was evolving in-tandem with another, we subjected our protein-coding gene sequences to the BGM algorithm which leverages Bayesian graphical models [51]. The BGM algorithm infers substitution history through the use of maximum-likelihood analyses for ancestral sequences and maps these to the phylogenetic tree, which allows for the detection of correlated patterns of substitution [51]. For our BGM analysis, we find evidence for 23 pairs of coevolving sites. This suggests interaction dynamics in tertiary space of the 3D, folded, protein level (see relevant sites in Figure 2) BDNF protein structure. Or, otherwise, is evidence for coevolving sites due to other fitness consequences related to compensation for maladaptive changes in one part of the protein sequence that may have occurred. When we review these sites, we notice that several pairs (see Figure 2) occur within alignment sites which correspond to the Human BDNF coordinate system (Table 2). These include sites (89, 184), (94, 155), (103, 233), (135, 154). Of note, several other sites also display interesting geometric features including triangular relations: [(81, 93), (93, 98), (81, 98)], an acyclic graph network of site connections [(70, 74), (74, 94), (94, 155)] and [(25, 49), (49, 85), (49, 86)], and more complex double linked co-evolutionary sites: [(39, 103), (103, 233)], and triple linked co-evolutionary sites: [(30, 119), (33, 119), (91, 119)]. Additionally, three-dimensional reconstruction - here focusing on a specific heterodimer configuration of BDNF and NT-4 as an example of a spatial protein-protein interaction - highlights that coevolving sites as well as positively evolving sites are likely to have been “fine tuned” over time to help support BDNF’s cognate functionality (see Figure 3). Mapping our FEL purifying sites in a structural configuration was not shown due to the overwhelming nature of negative selection acting on BDNF within mammals.

**Figure 2.**
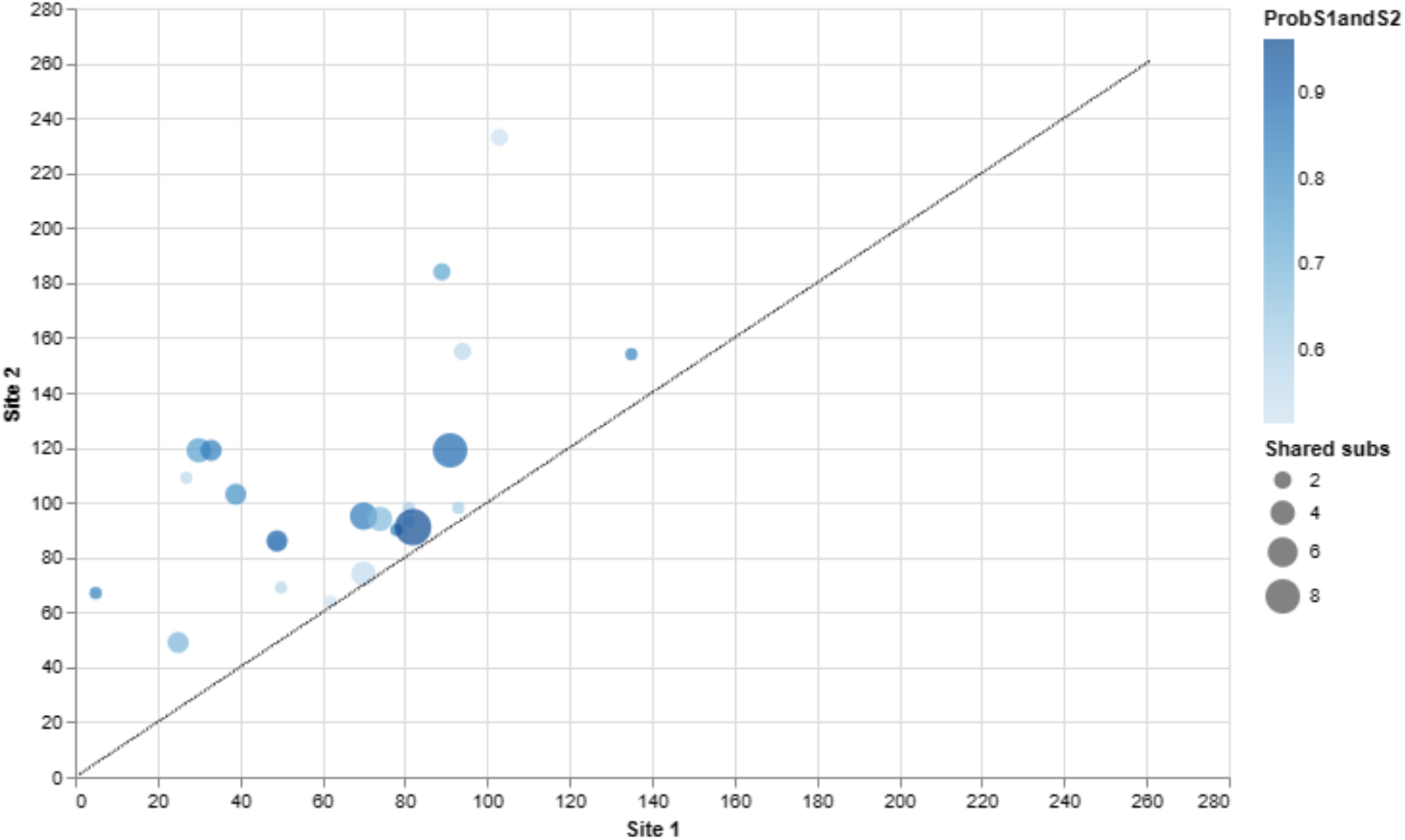
The BGM analysis of BDNF found 23 pairs of coevolving sites out of 261 total sites to be statistically significant (with a posterior probability threshold 0.5). Here, we plot only the statistically significant co-evolving pairs. The number of shared substitutions between pairs of co-evolving sites is visualized by the size of the circle, with larger circles indicating more shared substitutions. Poster probability of the interaction (coevolving pair) corresponds to the color blue, with dark blue indicating higher values. Individual BDNF sites are mapped on both the X and Y axis so that readers can view which sites are coevolving with another. Once more, note that the coevolution tends to be focally constrained to the broader BDNF prodomain region at a topological level, which is once more consistent with the idea that the NGF domain (site >144; see Figure 3) is highly conserved and probably deleteriously impacted by variation. However, we did discover four sites of coevolution in the NGF domain (basically, the mature BDNF protein) that are evolving with early prodomain sites. This highlights that both proximal and distal sites in BDNF can, and indeed are, evolving together over time.

**Figure 3.**
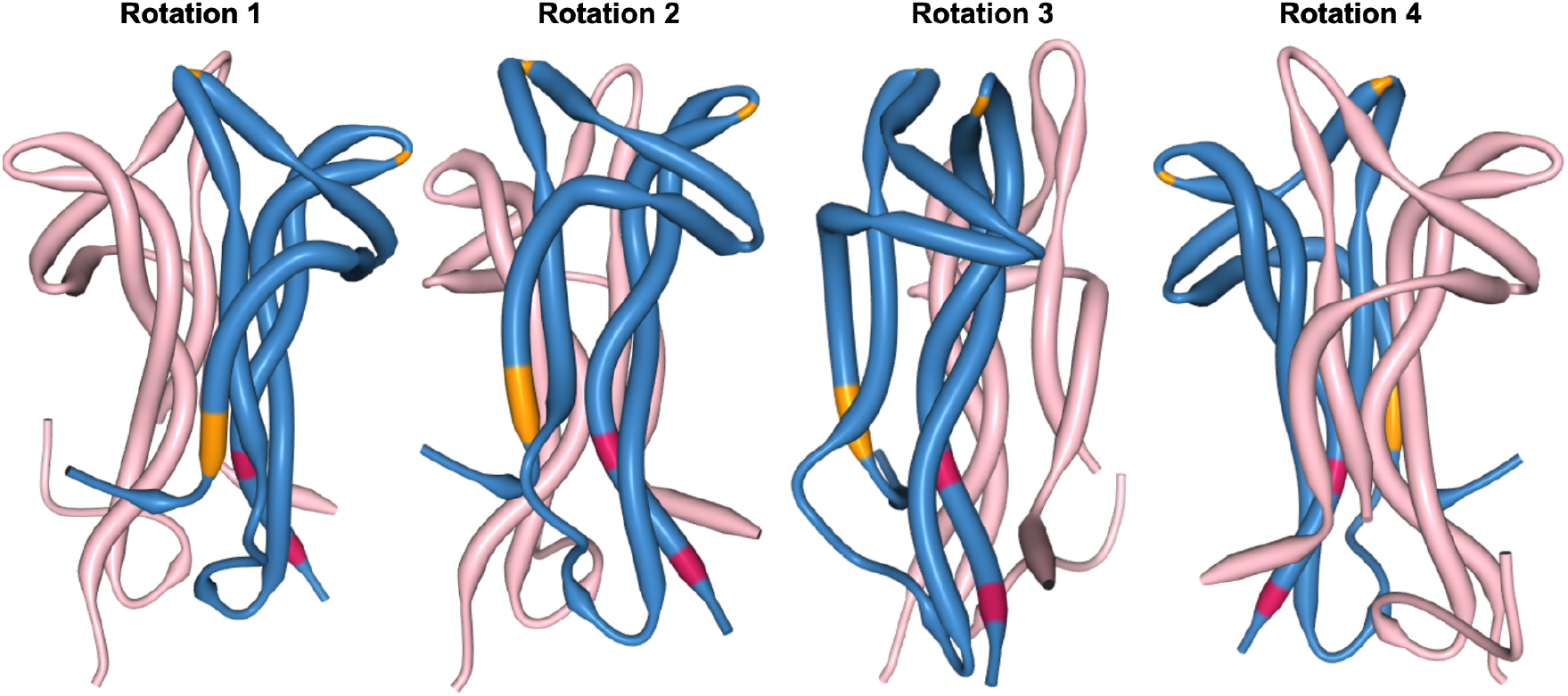
BDNF NT-4 Heterodimer Structural Analysis to Highlight Coevolving and Adaptive Sites at the 3D Protein Level. Demonstrating the structural configuration of the BDNF (blue) and NT-4 (pink) heterodimer (see https://www.rcsb.org/structure/1B8M), with rotations (arbitrary degrees) shown to accentuate view of coevolving sites (orange, see also Figure 2 and Table 3) and positively evolving sites (red; see Table 2). The PDB structure is limited largely to the NGF domain which limits our ability to highlight sites of interest (SOI), therefore we have limited our annotation only to the modeled sites in the structure. The relative positioning of coevolving and positively evolving sites in this heterodimer visualization (proximity to looping and other macro tertiary structures of protein). An interactive figure that is rotatable in 3D space, where occupations occur in three-dimensions (i.e. teasing out relative proximity in 2D linear space), is available here https://observablehq.com/@aglucaci/bdnf-structure.

**Table 3.** The BGM analysis of BDNF found 23 pairs of coevolving sites out of 261 total sites to be statistically significant (with a posterior probability threshold 0.5).

| # | Site 1 | Site 2 | P [Site 1 → Site 2] | P [Site 2 → Site 1] | P [Site 1 ↔ Site 2] | Site 1 subs | Site 2 subs | Shared subs |
| --- | --- | --- | --- | --- | --- | --- | --- | --- |
| 1 | 5 | 67 | 0.52 | 0.32 | 0.84 | 1 | 1 | 1 |
| 2 | 25 | 49 | 0.022 | 0.67 | 0.69 | 8 | 6 | 3 |
| 3 | 27 | 109 | 0.12 | 0.45 | 0.57 | 2 | 1 | 1 |
| 4 | 30 | 119 | 0.2 | 0.56 | 0.75 | 6 | 12 | 4 |
| 5 | 33 | 119 | 0.12 | 0.72 | 0.84 | 3 | 12 | 3 |
| 6 | 39 | 103 | 0.22 | 0.58 | 0.8 | 8 | 4 | 3 |
| 7 | 49 | 85 | 0.48 | 0.049 | 0.53 | 6 | 3 | 2 |
| 8 | 49 | 86 | 0.36 | 0.55 | 0.91 | 6 | 5 | 3 |
| 9 | 50 | 69 | 0.31 | 0.27 | 0.58 | 1 | 2 | 1 |
| 10 | 62 | 64 | 0.25 | 0.27 | 0.52 | 1 | 1 | 1 |
| 11 | 70 | 74 | 0.17 | 0.38 | 0.55 | 7 | 14 | 4 |
| 12 | 70 | 95 | 0.59 | 0.26 | 0.85 | 7 | 21 | 5 |
| 13 | 74 | 94 | 0.034 | 0.64 | 0.68 | 14 | 7 | 4 |
| 14 | 78 | 90 | 0.46 | 0.38 | 0.84 | 1 | 1 | 1 |
| 15 | 81 | 93 | 0.27 | 0.31 | 0.58 | 1 | 1 | 1 |
| 16 | 81 | 98 | 0.27 | 0.33 | 0.61 | 1 | 1 | 1 |
| 17 | 82 | 91 | 0.045 | 0.92 | 0.96 | 15 | 29 | 9 |
| 18 | 89 | 184 | 0.24 | 0.5 | 0.74 | 2 | 6 | 2 |
| 19 | 91 | 119 | 0.22 | 0.67 | 0.89 | 29 | 12 | 8 |
| 20 | 93 | 98 | 0.31 | 0.3 | 0.61 | 1 | 1 | 1 |
| 21 | 94 | 155 | 0.28 | 0.29 | 0.57 | 7 | 2 | 2 |
| 22 | 103 | 233 | 0.4 | 0.13 | 0.53 | 4 | 3 | 2 |
| 23 | 135 | 154 | 0.4 | 0.43 | 0.82 | 1 | 1 | 1 |

## DISCUSSION

In this study, we explore the evolutionary history of the BDNF gene in *Mammalia*. The BDNF gene is implicated in a number of human diseases including a variety of brain disorders such as neurodevelopmental disorders (e.g. autism spectrum disorder), neuropsychiatric disorders (e.g. depression, PTSD, schizophrenia), and some neurodegenerative disorders [1]. By using orthologous BDNF sequences within the *Mammalia* taxonomic group, our results indicate that within species, unique evolutionary pressures and site-specific changes within the BDNF gene have evolved across time. We performed a number of comparative evolutionary analyses to tease out signals from our orthologous gene collection in BDNF. Of note, the BDNF gene elicits tight regulation and specific functionality that can be separated from other neurotrophins, yet these growth factors remain closely related in their structure and sequence and conservation of the NGF functional domain. In the NGF domain, we observe a high degree of conservation (via purifying selection) across species, owing to the functional importance of this region in protein-protein interactions. This work additionally provides broad comparative insights into the evolutionary history of the BDNF gene family. Our MEME method identified novel substitutions (see Table 2) in regions of BDNF that may provide significant areas of interest for designing molecular therapeutic approaches, and their potential broader significance are outlined in further detail below.

### Predominant purifying selection across BDNF in Mammalia

Over time, evolution drives the divergence of genetic sequences. What can we learn from the direct comparison of the sequences of the BDNF gene in Mammalia? By comparing the BDNF products of orthologous genes in different species we observe the accumulation of mutations at different sites with varying degrees of insight into both BDNF functionality (see [29] for site annotation) and potential disease [1]. These are summarized in-full within Table S1 and Table 2. Coding sequences with highly constrained structures are expected to fix non-synonymous mutations at a slower rate due to the maladaptive nature of changes such as what we observe with the FEL negatively selected sites across BDNF. Additionally, we observe a high degree of negative (purifying) selection across the main functional domain (NGF) of BDNF. While structures for the NGF domain in most species under analysis do not exist, based on our findings we expect a highly conserved tertiary structure. Based on the high degree of purifying selection observed across BDNF, we hypothesize that BDNF plays a critical role in the underlying network of genes governing homeostasis and normal organismal development. This may have happened because BDNF is particularly useful specifically for the phylogenetic branch in question (i.e. mammals). This interpretation also tracks with the observation that BDNF is essential for normative development and is basically lethal in non-conditional full knock-out mammalian models.

### Non-Neutral Positive Diversifying Evolution Sites in the BDNF Gene

It has been described that BDNF particularly plays a role as a foundational gene for brain development [11]. Despite a significant level of purifying selection shaping the evolutionary history of BDNF (Figure 1), we observe several novel statistically significant sites under positive episodic diversifying selection across the BDNF gene (see Table 2). Traditionally, evolution of this variety consists of amino acid diversifying events that may promote phylogenetic adaptation and/or functionality. These results are entirely novel - they have not been previously reported (to the best of our knowledge) and MEME is an established and sensitive method for the analysis of episodic diversifying sites. Thus, the very specific and limited sites within BDNF to exhibit such patterns is a highly promising result from which to further disentangle BDNFs complex functionality and disease linkage. Presumably, we would encourage biologists to consider these sites as those that may contain important adaptive functions within the BDNF gene. However, where our results fall within the context of a core protein-protein interaction network of required genes for neural cellular diversity and development is yet to be determined. We do note that at least one identified site (238) overlaps with potential post-translational modifications to the human BDNF peptide (specifically, a disulfide bridge; see UniProt and [29]). This supports the idea that non-neutral positive diversifying sites within BDNF are not spurious and likely reflect specialized, regulatory, or functional capacities that may have yet to be annotated in-full. Given that this manuscript is devoted to analysis of BDNF’s evolution in mammals, we highlight the potential importance of these sites but emphasize that their importance remains a hypothesis that should be tested in well-defined experiments under controlled laboratory conditions.

### Discovery of Proximal & Distal Coevolving Site-Pairs in the BDNF Gene

Another novel, and potentially important, series of findings in this manuscript was the presence of numerous sites that exhibit coevolution. In fact, we observe a significant number of coevolving sites within the BDNF gene (see Figure 2, Table 3), and these too reflect an entirely novel aspect of BDNF biology that has not previously been reported. Evidence of coevolving sites are not limited to a particular domain (e.g. prodomain vs mature) nor specific motifs. Instead, coevolving pairs seem to be distributed across the entire BDNF gene with, perhaps unsurprisingly, an increased density of interactions early in the pre-pro domain region. However, we also note that there are coevolving sites in the mature NGF domain which are “linked” to early domain sites. Importantly, these relationships may confer strong epistatic interactions shaping the continued evolution of this critically important gene. The new evidence for coevolution may point to the importance of these sites in shaping the early regulatory or main functional (NGF) domain of BDNF. These residues may form important interactions for the functional integrity of BDNF and, importantly, the highly specific pairs which span the BDNF prodomain and its mature region point to a new mechanism by which the BDNF prodomain may have co-regulated the mature domain (or vice versa). Alternatively, these coevolving pairs may be part of a network of residues occupying a shifted fitness landscape in order to accommodate new or species-specific functional requirements.

### Potential Structural Implications of Evolving Sites

In considering our observation of both diversifying selection and coevolving sites in the BDNF gene, we considered the potential implications this may have at a protein structural level in three-dimensional space (see Figure 3). While protein structural impacts from evolution remain poorly understood and cannot be completely experimentally disentangled in a confirmatory sense, the implications fall upon our understanding of basic BDNF neurobiology. Here we note that our BGM and FEL analyses implicate the prodomain - the primary topological region of BDNF known for polymorphic variability (e.g. Val66Met, Gln75His) that is often linked to disease [1, 11, 29], and our 3D modeling suggests that two of our co-evolving sites appear to be associated with looping structures that could have important yet to be discovered functionality. In this regard, we predict that the evolutionary changes described here are likely to reflect some form of specialization and/or divergence in function and/or interaction partners at different points of BDNF’s evolutionary history in mammals. Thus, further work may unveil yet more novel sites that could provide further insight into the origins of BDNF’s diverse functionality and its role in disease.

### Limitations of our Computational Evolutionary Analysis

This analysis focused on BDNF sequences contained in the taxonomic group Mammalia in lieu of examining a more inclusive dataset for BDNF containing sequences from all of *Gnathostomata* (jawed vertebrates) or extension into invertebrate clades which may contain BDNF or BDNF-like analogue genes. Our results are applicable to mammals, which are our intentional taxonomic group of study, but we nonetheless emphasize that our results do not capture the *entirety* of BDNF’s evolutionary history (e.g. there could be more to learn about BDNF from birds, lizards, fish, and higher order taxonomic groups which we do not evaluate here). In addition we do not explore the patterns or mutational processes occurring outside of coding-sequence evolution which include complex structure and dynamics of non-coding regions in the BDNF gene or across. Therefore, evolutionary temporality is important in the context and interpretation of our results because Mammalia represents only a portion of the long evolutionary history of BDNF. Although we failed to find evidence for recombination in our dataset, species where we may find evidence for recombination may have been precluded from our analysis due to our decision to focus on Mammalian BDNF gene evolution. Further, a limitation of the current analysis is owed to the presence of indel events, especially in the early region of the alignment but which also occur in other spatially distributed regions of the BDNF gene. These indel events are not currently modeled in existing codon substitution models but may represent an additional pathway of evolutionary change. Nonetheless, the prominence of indels in our observations indicate that several regions of BDNF may evolve significantly through indel events across species. Lastly, although there is a risk that the “gappy” nature of the early region of our multiple sequence alignment may be a computational artifact of the alignment procedure, based on all other outputs we believe that our results are reasonably interpreted and have subsequently tolerated these potential effects.

### Future Directions: Understanding the Remainder of the Neurotrophin Family

We hypothesize that the similarities between neurotrophins reflects conserved evolutionary selection for motifs and domains for which support common functionality in neurotrophic factors between sites and lineages. While we note significant isotropy in mature peptide sequences for these factors, anisotropic pressures likely influenced the prodomain sequences of neurotrophins leading to alterations in processing, trafficking, regulation, and secretion. As such, we also predict differences in the evolutionary fate of other neurotrophins which also exhibit compartmentalized functionality due to similar alterations within their prodomains (i.e. similar results may be reasonably anticipated not just in BDNF, but also NGF, NT-3, and NT-4).

## CONCLUSION

To sum up, our research modeled the natural evolutionary history of changes in the BDNF gene across >160 mammalian genomes. Conservatively, this analysis spans approximately ∼177 million years of evolution - and going deeper could yet reveal more information on the ontogenesis of BDNF and its topological structure (and, consequently, function). Notably, we observed strict purifying selection in the main functional domain of the BDNF gene in mammals and discovered 6 specific sites in our homologous alignment that are under episodic selection in the early regulatory region of BDNF (i.e. the prodomain) and in the terminal region of BDNF. We also make the case for spatial coevolution within this gene, with 23 pair-sites that have evolved together. In sum, these data go above and beyond the common trope that “BDNF is highly conserved” by defining exactly where and how the mammalian BDNF has evolved. Thus, we confirm the widespread belief that the BDNF prodomain is more prone to change than the mature BDNF protein, having important implications for how we think about and consider genetic variation in BDNF and its linkage to disease.

## METHODS

### Data Retrieval

For this study, we queried the NCBI Ortholog database via https://www.ncbi.nlm.nih.gov/kis/ortholog/627/?scope=7776. For the purpose of this study, as we are interested in mammalian BDNF evolution, we limited our search to only include species from this taxonomic group (mammals, *Mammalia*). This returned 162 full gene transcripts and protein sequences. We downloaded all available files: RefSeq protein sequences, RefSeq transcript sequences, Tabular data (CSV, metadata). In Table 1, we provide a table of the species included in this analysis but we also make this accessible via GitHub. Furthermore, we also make these species NCBI accessions (see also Table 1) available for download on GitHub:

- AnalysisOfOrthologousCollections/BDNF_orthologs.csv at main · aglucaci/AnalysisOfOrthologousCollections · GitHub

### Data Cleaning

We used the protein sequence and full gene transcripts to derive coding sequences (CDS) (via a custom script, scripts/codons.py). However, this process was met with errors in 20 “PREDICTED” protein sequences which had invalid characters such as sequences which have incorrect ‘X’, or unresolved amino acids and these sequences were subsequently exempt from analysis. This process removes low-quality protein sequences from analysis which may inflate rates of nonsynonymous change.

Analysis of Orthologous Collections (AOC): Alignment, Recombination Detection, Tree inference & Selection Algorithms

The Analysis of Orthologous Collections (AOC) application is designed for comprehensive protein-coding molecular sequence analysis (https://github.com/aglucaci/AnalysisOfOrthologousCollections). It accomplishes this through a series of comparative evolutionary methods. AOC allows for the inclusion of recombination detection, a powerful force in shaping gene evolution and interpreting analytic results. As well, it allows for lineage assignment and annotation. This feature (lineage assignment) allows between group comparisons of selective pressures. This application currently accepts two input files: a protein sequence unaligned fasta file, and a transcript sequence unaligned fasta file for the same gene. Typically, this can be retrieved from public databases such as NCBI Orthologs. Although other methods of data compilation are also acceptable. In addition, the application is easily modifiable to accept a single CDS input, if that data is available.

If protein and transcript files are provided, a custom script ‘scripts/codons.py’ is executed and returns coding sequences where available. Note that this script currently is set to use the standard genetic code, this will need to be modified for alternate codon tables. This script also removes “low-quality” sequences if no match is found, see the above *Data cleaning* section.

**Step 1. Alignment.** We used the HyPhy [44] codon-aware multiple sequence alignment procedure available at (https://github.com/veg/hyphy-analyses/tree/master/codon-msa).

This was performed with a Human BDNF coding sequence *NM_001709.5 Homo sapiens brain derived neurotrophic factor (BDNF), transcript variant 4, mRNA* as a reference based alignment. Our alignment procedure retained 126 unique in-frame sequences.

**Step 2. Recombination detection.** Performed manually via RDP v5 [45], see below, the “Recombination detection” section for additional details. A recombination free file is placed in the following folder: results/BDNF/Recombinants. For the purpose of this study, we did not detect recombination in our dataset.

**Step 3. Tree inference and selection analyses.** For the recombination free fasta file, we perform maximum likelihood phylogenetic inference via IQ-TREE [46]. Next, the recombination free alignment and unrooted phylogenetic tree is evaluated through a standard suite of molecular evolutionary methods. This set of selection analyses include the following but for the sake of brevity some of these results were not shown (essentially, most were not statistically significant or not meaningful as relevant to the evolutionary results presented here).

- FEL: locates codon sites with evidence of pervasive positive diversifying or negative selection [47].
- BUSTEDS: tests for gene-wide episodic selection [48].
- MEME: locates codon sites with evidence of episodic positive diversifying selection, [49].
- aBSREL: tests if positive selection has occurred on a proportion of branches, [50].
- SLAC: performs substitution mapping, [47].
- BGM: identifies groups of sites that are apparently co-evolving, [51].
- RELAX: compare gene-wide selection pressure between the query clade and background sequences, [52].
- CFEL: comparison site-by-site selection pressure between query and background sequences, [53].
- FMM: examines model fit by permitting multiple instantaneous substitutions, [54].

**Step 4A.** Lineage assignment and tree annotation. For the unrooted phylogenetic tree, we perform lineage discovery, via NCBI and the python package ete3 toolkit. Assigning lineages to a K (by default, K = 20) number of taxonomic groups. Here, the aim is to have a broad representation of taxonomic groups, rather than the species being heavily clustered into a single group. As a reasonable approximation, we aim for <40% of species to be assigned to any one particular taxonomic group.

**Step 4B.** We perform tree labeling via the hyphy-analyses/Label-Trees (REF, link) method. Resulting in one annotated tree per lineage designation. For the purpose of this study, we will only consider the following five lineages for additional analyses (Artiodactyla, Carnivora, Chiroptera, Glires, Primates) as they are the most populated lineages.

**Step 5.** Selection analyses on lineages. Here, the recombination free fasta file and the set of annotated phylogenetic trees (where labeling was performed in Step 4) is provided for analysis with the RELAX and Contrast-FEL methods.

### Recombination detection

Manually tested via RDPv5.5 with modified settings as follows:

- We also included the following algorithms/analyses: RDP [55], GENECONV [56], Chimaera [57], MaxChi [58], BootScan [59] (Primary and Secondary Scan), SiScan [60] (Primary and Secondary Scan), 3Seq [61].
- Recombination events are ‘accepted’ in cases where 3 or more methods are in agreement.
- We slightly modified default parameters, such that

- Require topological evidence.
- Polish breakpoints.
- Check alignment consistency.
- Sequences are linear.
- List events detected by >2 methods.
- We manually recheck all of the events via “Recheck all identified events with all methods”.
- We manually accept events detected by >2 methods.
- The resulting alignment was saved as a distributed alignment (with recombinant regions separated).

Recombination was not detected within our Human reference based alignment. Therefore we used the single recombination free alignment for analyses.

### Data & Software Availability Statement

The AOC application is freely available via a dedicated GitHub repository at: https://github.com/aglucaci/AnalysisOfOrthologousCollections

Raw data for this study is available on GitHub: https://github.com/aglucaci/AnalysisOfOrthologousCollections/tree/main/data/BDNF

Full results for this study include all HyPhy selection analyses JSON-formatted result files are available on GitHub: https://github.com/aglucaci/AnalysisOfOrthologousCollections/tree/main/results/BDNF

## Contributions & Acknowledgements

M.N. and A.L. contributed equally. M.N. was supported by a NHMRC CJ Martin Fellowship for stem cell training at Weill Cornell Medical College.

## Supplementary Material

**Table S1.** The FEL analysis of the BDNF gene found 174 of 261 (66.7%) sites to be statistically significant (LRT p-value <= 0.1) for pervasive negative (purifying) selection.

| Site | alpha | beta | alpha=beta | LRT | p-value | Total branch length | dN/dS LB | dN/dS MLE | dN/dS UB |
| --- | --- | --- | --- | --- | --- | --- | --- | --- | --- |
| 1 | 0.0000 | 0.0000 | 0.0000 | 0.0000 | 1.0000 | 0.0000 | 0.0000 | 0.0000 | 0.0000 |
| 2 | 0.0000 | 0.0000 | 0.0000 | 0.0000 | 1.0000 | 0.0000 | 0.0000 | 0.0000 | 0.0000 |
| 3 | 0.0000 | 0.0000 | 0.0000 | 0.0000 | 1.0000 | 0.0000 | 0.0000 | 0.0000 | 0.0000 |
| 4 | 0.1116 | 0.0635 | 0.0811 | 0.1573 | 0.6916 | 0.6594 | 0.0325 | 0.5690 | 2.5205 |
| 5 | 0.0000 | 0.0564 | 0.0468 | 0.3606 | 0.5482 | 0.3806 | 1462.7972 | 10000.0000 | 10000.0000 |
| 6 | 0.2221 | 0.0000 | 0.0805 | 4.0430 | 0.0444 | 0.6545 | 0.0000 | 0.0000 | 0.5478 |
| 7 | 0.0000 | 0.0000 | 0.0000 | 0.0000 | 1.0000 | 0.0000 | 0.0000 | 0.0000 | 0.0000 |
| 8 | 0.0000 | 0.0000 | 0.0000 | 0.0000 | 1.0000 | 0.0000 | 0.0000 | 0.0000 | 0.0000 |
| 9 | 0.0928 | 0.0000 | 0.0397 | 2.0198 | 0.1553 | 0.3227 | 0.0000 | 0.0000 | 1.2848 |
| 10 | 0.1594 | 0.0000 | 0.0409 | 2.7359 | 0.0981 | 0.3326 | 0.0000 | 0.0000 | 0.6580 |
| 11 | 0.1729 | 0.0000 | 0.0900 | 3.1185 | 0.0774 | 0.7317 | 0.0000 | 0.0000 | 0.9176 |
| 12 | 0.0000 | 0.0000 | 0.0000 | 0.0000 | 1.0000 | 0.0000 | 0.0000 | 0.0000 | 0.0000 |
| 13 | 1.9630 | 0.0000 | 0.2850 | 22.6368 | 0.0000 | 2.3174 | 0.0000 | 0.0000 | 0.0549 |
| 14 | 0.0000 | 0.2172 | 0.1438 | 3.0861 | 0.0790 | 1.1694 | 6185.5906 | 10000.000<br>0 | 10000.000<br>0 |
| 15 | 0.0000 | 0.0000 | 0.0000 | 0.0000 | 1.0000 | 0.0000 | 0.0000 | 0.0000 | 0.0000 |
| 16 | 0.0000 | 0.0000 | 0.0000 | 0.0000 | 1.0000 | 0.0000 | 0.0000 | 0.0000 | 0.0000 |
| 17 | 0.2673 | 0.0530 | 0.0885 | 1.1690 | 0.2796 | 0.7199 | 0.0113 | 0.1982 | 0.8804 |
| 18 | 0.3413 | 0.0000 | 0.1102 | 6.7673 | 0.0093 | 0.8958 | 0.0000 | 0.0000 | 0.3054 |
| 19 | 0.4344 | 0.0000 | 0.1182 | 7.7643 | 0.0053 | 0.9612 | 0.0000 | 0.0000 | 0.2399 |
| 20 | 0.7437 | 0.0000 | 0.1993 | 12.7922 | 0.0003 | 1.6204 | 0.0000 | 0.0000 | 0.1425 |
| 21 | 0.0000 | 0.0000 | 0.0000 | 0.0000 | 1.0000 | 0.0000 | 0.0000 | 0.0000 | 0.0000 |
| 22 | 0.7734 | 0.0000 | 0.1935 | 13.6496 | 0.0002 | 1.5734 | 0.0000 | 0.0000 | 0.1283 |
| 23 | 0.1435 | 0.0527 | 0.0773 | 0.4750 | 0.4907 | 0.6284 | 0.0210 | 0.3670 | 1.6325 |
| 24 | 1.7963 | 0.4672 | 0.8629 | 8.8413 | 0.0029 | 7.0170 | 0.1183 | 0.2601 | 0.4893 |
| 25 | 1.4891 | 0.3979 | 0.5537 | 4.2623 | 0.0390 | 4.5023 | 0.1213 | 0.2672 | 0.5042 |
| 26 | 0.1396 | 1.0896 | 0.7817 | 6.8872 | 0.0087 | 6.3560 | 4.5038 | 7.8049 | 12.5990 |
| 27 | 0.2849 | 0.2004 | 0.2436 | 0.1449 | 0.7035 | 1.9806 | 0.1198 | 0.7033 | 2.1145 |
| 28 | 0.0000 | 0.0000 | 0.0000 | 0.0000 | 1.0000 | 0.0000 | 0.0000 | 0.0000 | 0.0000 |
| 29 | 0.4139 | 0.0566 | 0.1834 | 4.1399 | 0.0419 | 1.4914 | 0.0078 | 0.1367 | 0.6064 |
| 30 | 0.0000 | 0.3797 | 0.2579 | 4.6268 | 0.0315 | 2.0973 | 7260.0520 | 10000.000<br>0 | 10000.000<br>0 |
| 31 | 0.2774 | 0.0000 | 0.0763 | 5.1487 | 0.0233 | 0.6206 | 0.0000 | 0.0000 | 0.3653 |
| 32 | 0.2572 | 0.4242 | 0.3976 | 0.2465 | 0.6196 | 3.2327 | 0.7845 | 1.6493 | 3.0082 |
| 33 | 0.3030 | 0.2456 | 0.2654 | 0.0484 | 0.8259 | 2.1581 | 0.2009 | 0.8105 | 2.1234 |
| 34 | 1.0536 | 0.1089 | 0.3602 | 9.9540 | 0.0016 | 2.9291 | 0.0172 | 0.1034 | 0.3222 |
| 35 | 0.0000 | 0.0000 | 0.0000 | 0.0000 | 1.0000 | 0.0000 | 0.0000 | 0.0000 | 0.0000 |
| 36 | 0.2906 | 0.0686 | 0.1107 | 0.9484 | 0.3301 | 0.9000 | 0.0134 | 0.2360 | 1.0454 |
| 37 | 0.8023 | 0.0000 | 0.2613 | 15.6626 | 0.0001 | 2.1248 | 0.0000 | 0.0000 | 0.1321 |
| 38 | 0.7113 | 0.3303 | 0.4528 | 1.6696 | 0.1963 | 3.6817 | 0.1837 | 0.4644 | 0.9547 |
| 39 | 0.4421 | 0.5250 | 0.5030 | 0.0485 | 0.8256 | 4.0903 | 0.5420 | 1.1877 | 2.2205 |
| 40 | 0.4097 | 0.0000 | 0.1578 | 7.5858 | 0.0059 | 1.2834 | 0.0000 | 0.0000 | 0.3013 |
| 41 | 0.0000 | 0.0551 | 0.0393 | 0.6774 | 0.4105 | 0.3195 | 1462.7591 | 10000.000<br>0 | 10000.000<br>0 |
| 42 | 1.2391 | 0.0506 | 0.2862 | 13.8667 | 0.0002 | 2.3274 | 0.0023 | 0.0409 | 0.1816 |
| 43 | 0.1559 | 0.0000 | 0.0396 | 2.7457 | 0.0975 | 0.3218 | 0.0000 | 0.0000 | 0.6535 |
| 44 | 0.3404 | 0.0556 | 0.1494 | 2.9408 | 0.0864 | 1.2148 | 0.0093 | 0.1634 | 0.7237 |
| 45 | 0.0928 | 0.0000 | 0.0443 | 1.4933 | 0.2217 | 0.3605 | 0.0000 | 0.0000 | 1.7490 |
| 46 | 0.0000 | 0.1640 | 0.1370 | 1.0523 | 0.3050 | 1.1140 | 5270.0970 | 10000.000<br>0 | 10000.000<br>0 |
| 47 | 1.8271 | 0.1373 | 0.3286 | 11.8421 | 0.0006 | 2.6721 | 0.0187 | 0.0751 | 0.1965 |
| 48 | 0.7118 | 1.0257 | 0.9372 | 0.4209 | 0.5165 | 7.6211 | 0.8165 | 1.4411 | 2.3527 |
| 49 | 1.3089 | 0.3744 | 0.5850 | 4.9417 | 0.0262 | 4.7570 | 0.1222 | 0.2860 | 0.5605 |
| 50 | 0.2882 | 0.0532 | 0.1161 | 2.0383 | 0.1534 | 0.9440 | 0.0105 | 0.1844 | 0.8138 |
| 51 | 0.1419 | 0.0000 | 0.0395 | 2.5243 | 0.1121 | 0.3214 | 0.0000 | 0.0000 | 0.7470 |
| 52 | 0.2736 | 0.3297 | 0.3204 | 0.0312 | 0.8599 | 2.6057 | 0.4762 | 1.2050 | 2.4754 |
| 53 | 1.5426 | 0.4637 | 0.8202 | 7.0701 | 0.0078 | 6.6692 | 0.1365 | 0.3006 | 0.5668 |
| 54 | 1.5617 | 0.2216 | 0.6252 | 13.3081 | 0.0003 | 5.0838 | 0.0440 | 0.1419 | 0.3335 |
| 55 | 3.6302 | 0.5463 | 1.6240 | 22.5001 | 0.0000 | 13.2053 | 0.0643 | 0.1505 | 0.2957 |
| 56 | 1.5021 | 0.0545 | 0.4719 | 20.6584 | 0.0000 | 3.8374 | 0.0021 | 0.0363 | 0.1612 |
| 57 | 0.5526 | 0.2782 | 0.3576 | 0.9810 | 0.3220 | 2.9080 | 0.1797 | 0.5034 | 1.0950 |
| 58 | 0.3979 | 0.0869 | 0.2313 | 2.3382 | 0.1262 | 1.8808 | 0.0124 | 0.2184 | 0.9655 |
| 59 | 11.1190 | 0.8319 | 1.4313 | 1.5693 | 0.2103 | 11.6383 | 0.0038 | 0.0748 | 0.6522 |
| 60 | 4.3720 | 0.3686 | 1.3037 | 33.3567 | 0.0000 | 10.6014 | 0.0332 | 0.0843 | 0.1727 |
| 61 | 0.0000 | 0.0000 | 0.0000 | 0.0000 | 1.0000 | 0.0000 | 0.0000 | 0.0000 | 0.0000 |
| 62 | 1.7721 | 2193.2857 | 2856.9387 | 0.0000 | 1.0000 | 23231.3076 | 0.3292 | 1237.7020 | 10000.0000 |
| 63 | 0.0000 | 3.6440 | 2.0642 | 0.9793 | 0.3224 | 16.7849 | 1464.9793 | 10000.0000 | 10000.0000 |
| 64 | 7.4868 | 0.0000 | 1.0937 | 6.3427 | 0.0118 | 8.8931 | 0.0000 | 0.0000 | 0.1725 |
| 65 | 122.5088 | 0.0000 | 1.1740 | 7.2662 | 0.0070 | 9.5464 | 0.0000 | 0.0000 | 0.0108 |
| 66 | 2.8095 | 0.0000 | 0.7275 | 4.7774 | 0.0288 | 5.9161 | 0.0000 | 0.0000 | 0.3619 |
| 67 | 10.8121 | 0.0000 | 1.6286 | 13.5042 | 0.0002 | 13.2432 | 0.0000 | 0.0000 | 0.0777 |
| 68 | 4.1586 | 0.4872 | 1.5564 | 8.0811 | 0.0045 | 12.6556 | 0.0193 | 0.1172 | 0.3731 |
| 69 | 3.2764 | 0.3962 | 1.3489 | 22.0059 | 0.0000 | 10.9685 | 0.0431 | 0.1209 | 0.2630 |
| 70 | 0.9289 | 0.0000 | 0.3524 | 13.2397 | 0.0003 | 2.8657 | 0.0000 | 0.0000 | 0.1693 |
| 71 | 1.1053 | 0.0000 | 0.3383 | 20.8817 | 0.0000 | 2.7513 | 0.0000 | 0.0000 | 0.0943 |
| 72 | 0.2921 | 0.0985 | 0.1261 | 0.6630 | 0.4155 | 1.0251 | 0.0558 | 0.3373 | 1.0461 |
| 73 | 1.1053 | 0.0000 | 0.3395 | 20.5444 | 0.0000 | 2.7603 | 0.0000 | 0.0000 | 0.0954 |
| 74 | 2.9165 | 0.0575 | 0.6252 | 32.3768 | 0.0000 | 5.0841 | 0.0011 | 0.0197 | 0.0877 |
| 75 | 0.8228 | 0.0000 | 0.2007 | 13.9151 | 0.0002 | 1.6320 | 0.0000 | 0.0000 | 0.1241 |
| 76 | 5.5185 | 0.0000 | 0.5700 | 53.7705 | 0.0000 | 4.6353 | 0.0000 | 0.0000 | 0.0174 |
| 77 | 0.6280 | 0.0619 | 0.2215 | 5.6574 | 0.0174 | 1.8010 | 0.0056 | 0.0985 | 0.4366 |
| 78 | 2.1641 | 0.0489 | 0.3787 | 23.0125 | 0.0000 | 3.0792 | 0.0013 | 0.0226 | 0.1005 |
| 79 | 1.8388 | 0.0000 | 0.4162 | 29.0685 | 0.0000 | 3.3840 | 0.0000 | 0.0000 | 0.0556 |
| 80 | 0.2839 | 0.0541 | 0.0909 | 1.2074 | 0.2718 | 0.7393 | 0.0109 | 0.1905 | 0.8475 |
| 81 | 1.3529 | 0.0000 | 0.5463 | 20.2275 | 0.0000 | 4.4420 | 0.0000 | 0.0000 | 0.1126 |
| 82 | 1.8857 | 0.0830 | 0.7606 | 19.4503 | 0.0000 | 6.1851 | 0.0025 | 0.0440 | 0.1950 |
| 83 | 2.4273 | 0.1547 | 0.5523 | 20.6750 | 0.0000 | 4.4910 | 0.0157 | 0.0637 | 0.1663 |
| 84 | 0.5934 | 0.0000 | 0.0911 | 7.3673 | 0.0066 | 0.7411 | 0.0000 | 0.0000 | 0.1756 |
| 85 | 0.3406 | 0.1489 | 0.1729 | 0.4375 | 0.5083 | 1.4059 | 0.1081 | 0.4371 | 1.1399 |
| 86 | 0.0000 | 0.3235 | 0.2567 | 2.3672 | 0.1239 | 2.0875 | 6809.7393 | 10000.000<br>0 | 10000.000<br>0 |
| 87 | 1.2767 | 0.1562 | 0.4047 | 9.9233 | 0.0016 | 3.2907 | 0.0303 | 0.1223 | 0.3201 |
| 88 | 1.3410 | 0.5707 | 0.8242 | 3.2658 | 0.0707 | 6.7019 | 0.2024 | 0.4256 | 0.7810 |
| 89 | 1.2511 | 0.1289 | 0.5092 | 12.0135 | 0.0005 | 4.1408 | 0.0170 | 0.1031 | 0.3194 |
| 90 | 0.6067 | 0.0552 | 0.2020 | 6.0849 | 0.0136 | 1.6429 | 0.0052 | 0.0910 | 0.4030 |
| 91 | 4.6374 | 1.7289 | 2.1855 | 6.6899 | 0.0097 | 17.7717 | 0.2477 | 0.3728 | 0.5401 |
| 92 | 1.8962 | 0.0535 | 0.4607 | 23.2267 | 0.0000 | 3.7463 | 0.0016 | 0.0282 | 0.1245 |
| 93 | 3.5662 | 0.0542 | 0.7407 | 40.9892 | 0.0000 | 6.0226 | 0.0009 | 0.0152 | 0.0676 |
| 94 | 1.3252 | 0.3547 | 0.4855 | 3.6636 | 0.0556 | 3.9476 | 0.1139 | 0.2676 | 0.5226 |
| 95 | 1.3323 | 1.5991 | 1.5317 | 0.1473 | 0.7011 | 12.4555 | 0.7548 | 1.2003 | 1.8159 |
| 96 | 1.3529 | 0.0000 | 0.1787 | 14.6186 | 0.0001 | 1.4535 | 0.0000 | 0.0000 | 0.0749 |
| 97 | 1.8434 | 0.0489 | 0.2953 | 17.5494 | 0.0000 | 2.4011 | 0.0015 | 0.0265 | 0.1178 |
| 98 | 0.6510 | 0.0536 | 0.2046 | 6.5994 | 0.0102 | 1.6636 | 0.0047 | 0.0824 | 0.3692 |
| 99 | 0.0000 | 0.0000 | 0.0000 | 0.0000 | 1.0000 | 0.0000 | 0.0000 | 0.0000 | 0.0000 |
| 100 | 0.5931 | 0.3419 | 0.4207 | 0.4996 | 0.4797 | 3.4205 | 0.1807 | 0.5764 | 1.3251 |
| 101 | 5.6241 | 0.0678 | 0.7121 | 36.3681 | 0.0000 | 5.7902 | 0.0007 | 0.0120 | 0.0536 |
| 102 | 2.4986 | 0.1708 | 0.7664 | 25.4012 | 0.0000 | 6.2322 | 0.0169 | 0.0684 | 0.1784 |
| 103 | 0.6960 | 0.0000 | 0.2116 | 11.7725 | 0.0006 | 1.7204 | 0.0000 | 0.0000 | 0.1687 |
| 104 | 2.3048 | 0.0000 | 0.5043 | 34.9881 | 0.0000 | 4.1010 | 0.0000 | 0.0000 | 0.0444 |
| 105 | 0.6641 | 0.0000 | 0.1786 | 10.1702 | 0.0014 | 1.4520 | 0.0000 | 0.0000 | 0.1795 |
| 106 | 0.0000 | 0.0000 | 0.0000 | 0.0000 | 1.0000 | 0.0000 | 0.0000 | 0.0000 | 0.0000 |
| 107 | 0.9618 | 0.0000 | 0.3094 | 15.6026 | 0.0001 | 2.5158 | 0.0000 | 0.0000 | 0.1311 |
| 108 | 1.7507 | 0.0000 | 0.2683 | 25.8194 | 0.0000 | 2.1813 | 0.0000 | 0.0000 | 0.0495 |
| 109 | 1.2561 | 0.0472 | 0.2636 | 13.7621 | 0.0002 | 2.1434 | 0.0021 | 0.0376 | 0.1661 |
| 110 | 0.3333 | 0.0000 | 0.0900 | 5.1186 | 0.0237 | 0.7317 | 0.0000 | 0.0000 | 0.3586 |
| 111 | 0.0000 | 0.0000 | 0.0000 | 0.0000 | 1.0000 | 0.0000 | 0.0000 | 0.0000 | 0.0000 |
| 112 | 0.2221 | 0.0000 | 0.0736 | 4.3993 | 0.0360 | 0.5984 | 0.0000 | 0.0000 | 0.4774 |
| 113 | 0.2700 | 0.0000 | 0.1012 | 3.9075 | 0.0481 | 0.8232 | 0.0000 | 0.0000 | 0.5785 |
| 114 | 0.6015 | 0.0000 | 0.0925 | 7.5591 | 0.0060 | 0.7518 | 0.0000 | 0.0000 | 0.1729 |
| 115 | 2.1843 | 0.0000 | 0.5950 | 37.4897 | 0.0000 | 4.8380 | 0.0000 | 0.0000 | 0.0485 |
| 116 | 1.7115 | 0.0000 | 0.4751 | 29.9207 | 0.0000 | 3.8634 | 0.0000 | 0.0000 | 0.0619 |
| 117 | 1.3529 | 0.0000 | 0.4841 | 22.7688 | 0.0000 | 3.9361 | 0.0000 | 0.0000 | 0.0965 |
| 118 | 0.2799 | 0.0000 | 0.0871 | 4.6517 | 0.0310 | 0.7081 | 0.0000 | 0.0000 | 0.4354 |
| 119 | 2.6166 | 0.0000 | 0.5451 | 36.1605 | 0.0000 | 4.4325 | 0.0000 | 0.0000 | 0.0423 |
| 120 | 0.0928 | 0.0000 | 0.0445 | 1.5555 | 0.2123 | 0.3619 | 0.0000 | 0.0000 | 1.7079 |
| 121 | 3.0538 | 0.0000 | 0.9175 | 44.7049 | 0.0000 | 7.4605 | 0.0000 | 0.0000 | 0.0424 |
| 122 | 0.0000 | 0.0000 | 0.0000 | 0.0000 | 1.0000 | 0.0000 | 0.0000 | 0.0000 | 0.0000 |
| 123 | 0.5018 | 0.0000 | 0.1210 | 8.4758 | 0.0036 | 0.9840 | 0.0000 | 0.0000 | 0.2034 |
| 124 | 0.0000 | 0.0000 | 0.0000 | 0.0000 | 1.0000 | 0.0000 | 0.0000 | 0.0000 | 0.0000 |
| 125 | 0.0000 | 0.0000 | 0.0000 | 0.0000 | 1.0000 | 0.0000 | 0.0000 | 0.0000 | 0.0000 |
| 126 | 0.1674 | 0.0519 | 0.0785 | 0.6645 | 0.4150 | 0.6381 | 0.0174 | 0.3101 | 1.3547 |
| 127 | 0.5910 | 0.0680 | 0.1651 | 3.2197 | 0.0728 | 1.3427 | 0.0066 | 0.1151 | 0.5099 |
| 128 | 0.4829 | 0.0000 | 0.2240 | 7.5898 | 0.0059 | 1.8217 | 0.0000 | 0.0000 | 0.3344 |
| 129 | 0.7281 | 0.0000 | 0.1574 | 12.1458 | 0.0005 | 1.2801 | 0.0000 | 0.0000 | 0.1324 |
| 130 | 1.3529 | 0.0000 | 0.4223 | 26.4290 | 0.0000 | 3.4343 | 0.0000 | 0.0000 | 0.0770 |
| 131 | 1.3529 | 0.0000 | 0.4072 | 24.6703 | 0.0000 | 3.3113 | 0.0000 | 0.0000 | 0.0770 |
| 132 | 0.5994 | 0.0000 | 0.0843 | 7.7423 | 0.0054 | 0.6853 | 0.0000 | 0.0000 | 0.1579 |
| 133 | 0.0000 | 0.0000 | 0.0000 | 0.0000 | 1.0000 | 0.0000 | 0.0000 | 0.0000 | 0.0000 |
| 134 | 2.4575 | 0.0000 | 0.6712 | 37.7462 | 0.0000 | 5.4579 | 0.0000 | 0.0000 | 0.0483 |
| 135 | 0.0000 | 0.0435 | 0.0438 | 0.0008 | 0.9774 | 0.3558 | 1462.1672 | 10000.000<br>0 | 10000.000<br>0 |
| 136 | 1.0696 | 0.0000 | 0.2395 | 17.4870 | 0.0000 | 1.9473 | 0.0000 | 0.0000 | 0.0925 |
| 137 | 0.4335 | 0.0000 | 0.1301 | 7.1875 | 0.0073 | 1.0578 | 0.0000 | 0.0000 | 0.2750 |
| 138 | 0.6509 | 0.0000 | 0.2392 | 11.8685 | 0.0006 | 1.9451 | 0.0000 | 0.0000 | 0.1867 |
| 139 | 0.4217 | 0.0000 | 0.1161 | 7.6814 | 0.0056 | 0.9442 | 0.0000 | 0.0000 | 0.2443 |
| 140 | 0.9836 | 0.0000 | 0.1277 | 11.8927 | 0.0006 | 1.0384 | 0.0000 | 0.0000 | 0.0967 |
| 141 | 3.5104 | 0.0000 | 0.9082 | 50.5522 | 0.0000 | 7.3849 | 0.0000 | 0.0000 | 0.0344 |
| 142 | 0.0000 | 0.0000 | 0.0000 | 0.0000 | 1.0000 | 0.0000 | 0.0000 | 0.0000 | 0.0000 |
| 143 | 3.8890 | 0.0000 | 0.8487 | 57.0286 | 0.0000 | 6.9012 | 0.0000 | 0.0000 | 0.0273 |
| 144 | 0.5803 | 0.0000 | 0.1576 | 10.3294 | 0.0013 | 1.2818 | 0.0000 | 0.0000 | 0.1796 |
| 145 | 0.8551 | 0.0000 | 0.2336 | 15.3582 | 0.0001 | 1.8993 | 0.0000 | 0.0000 | 0.1209 |
| 146 | 3.1941 | 0.0000 | 0.8296 | 52.1403 | 0.0000 | 6.7458 | 0.0000 | 0.0000 | 0.0335 |
| 147 | 0.4576 | 0.0000 | 0.1172 | 8.0791 | 0.0045 | 0.9527 | 0.0000 | 0.0000 | 0.2215 |
| 148 | 0.0000 | 0.0000 | 0.0000 | 0.0000 | 1.0000 | 0.0000 | 0.0000 | 0.0000 | 0.0000 |
| 149 | 0.1879 | 0.0000 | 0.0883 | 2.9817 | 0.0842 | 0.7177 | 0.0000 | 0.0000 | 0.8603 |
| 150 | 2.4574 | 0.0000 | 0.2729 | 29.1158 | 0.0000 | 2.2192 | 0.0000 | 0.0000 | 0.0347 |
| 151 | 1.3529 | 0.0000 | 0.3641 | 20.8617 | 0.0000 | 2.9610 | 0.0000 | 0.0000 | 0.0881 |
| 152 | 1.8431 | 0.0000 | 0.3369 | 26.2702 | 0.0000 | 2.7398 | 0.0000 | 0.0000 | 0.0543 |
| 153 | 0.0000 | 0.0000 | 0.0000 | 0.0000 | 1.0000 | 0.0000 | 0.0000 | 0.0000 | 0.0000 |
| 154 | 1.1326 | 0.0449 | 0.2254 | 12.6183 | 0.0004 | 1.8326 | 0.0023 | 0.0397 | 0.1759 |
| 155 | 1.9612 | 0.1105 | 0.5215 | 19.7057 | 0.0000 | 4.2410 | 0.0093 | 0.0564 | 0.1754 |
| 156 | 9.5442 | 0.0000 | 0.6185 | 72.6680 | 0.0000 | 5.0296 | 0.0000 | 0.0000 | 0.0090 |
| 157 | 0.5748 | 0.0547 | 0.1375 | 3.7909 | 0.0515 | 1.1182 | 0.0054 | 0.0951 | 0.4215 |
| 158 | 0.0000 | 0.0000 | 0.0000 | 0.0000 | 1.0000 | 0.0000 | 0.0000 | 0.0000 | 0.0000 |
| 159 | 2.8095 | 0.0000 | 0.8095 | 44.3396 | 0.0000 | 6.5824 | 0.0000 | 0.0000 | 0.0424 |
| 160 | 2.4575 | 0.0000 | 0.6733 | 41.9754 | 0.0000 | 5.4754 | 0.0000 | 0.0000 | 0.0429 |
| 161 | 3.1758 | 0.0000 | 0.7134 | 48.2008 | 0.0000 | 5.8007 | 0.0000 | 0.0000 | 0.0328 |
| 162 | 1.3529 | 0.0000 | 0.3849 | 24.2341 | 0.0000 | 3.1298 | 0.0000 | 0.0000 | 0.0770 |
| 163 | 1.6026 | 0.0501 | 0.3633 | 19.2676 | 0.0000 | 2.9542 | 0.0018 | 0.0313 | 0.1388 |
| 164 | 0.6094 | 0.0000 | 0.1545 | 10.8904 | 0.0010 | 1.2565 | 0.0000 | 0.0000 | 0.1629 |
| 165 | 0.2724 | 0.0000 | 0.0441 | 3.6202 | 0.0571 | 0.3588 | 0.0000 | 0.0000 | 0.3729 |
| 166 | 1.3529 | 0.0000 | 0.3828 | 23.3448 | 0.0000 | 3.1125 | 0.0000 | 0.0000 | 0.0779 |
| 167 | 1.3492 | 0.0541 | 0.4779 | 20.1541 | 0.0000 | 3.8862 | 0.0023 | 0.0401 | 0.1781 |
| 168 | 0.0000 | 0.0000 | 0.0000 | 0.0000 | 1.0000 | 0.0000 | 0.0000 | 0.0000 | 0.0000 |
| 169 | 0.0000 | 0.0000 | 0.0000 | 0.0000 | 1.0000 | 0.0000 | 0.0000 | 0.0000 | 0.0000 |
| 170 | 0.0000 | 0.0000 | 0.0000 | 0.0000 | 1.0000 | 0.0000 | 0.0000 | 0.0000 | 0.0000 |
| 171 | 2.3512 | 0.0000 | 0.6698 | 30.8621 | 0.0000 | 5.4463 | 0.0000 | 0.0000 | 0.0603 |
| 172 | 1.3529 | 0.0000 | 0.3510 | 23.6433 | 0.0000 | 2.8544 | 0.0000 | 0.0000 | 0.0750 |
| 173 | 0.9159 | 0.0000 | 0.2362 | 16.1752 | 0.0001 | 1.9207 | 0.0000 | 0.0000 | 0.1110 |
| 174 | 2.5514 | 0.0000 | 0.5850 | 39.0254 | 0.0000 | 4.7569 | 0.0000 | 0.0000 | 0.0413 |
| 175 | 0.4238 | 0.0000 | 0.1290 | 7.0778 | 0.0078 | 1.0492 | 0.0000 | 0.0000 | 0.2813 |
| 176 | 3.0000 | 0.0000 | 0.8020 | 50.0929 | 0.0000 | 6.5213 | 0.0000 | 0.0000 | 0.0351 |
| 177 | 0.2903 | 0.0000 | 0.0867 | 4.8037 | 0.0284 | 0.7050 | 0.0000 | 0.0000 | 0.4106 |
| 178 | 2.0769 | 0.0000 | 0.7435 | 33.6840 | 0.0000 | 6.0457 | 0.0000 | 0.0000 | 0.0637 |
| 179 | 1.3529 | 0.0000 | 0.2916 | 20.9924 | 0.0000 | 2.3712 | 0.0000 | 0.0000 | 0.0756 |
| 180 | 2.8095 | 0.0000 | 0.5579 | 41.9086 | 0.0000 | 4.5370 | 0.0000 | 0.0000 | 0.0356 |
| 181 | 0.4269 | 0.0000 | 0.1294 | 7.1083 | 0.0077 | 1.0522 | 0.0000 | 0.0000 | 0.2792 |
| 182 | 2.0769 | 0.0000 | 0.5715 | 36.1609 | 0.0000 | 4.6469 | 0.0000 | 0.0000 | 0.0510 |
| 183 | 2.4177 | 0.0000 | 0.7387 | 39.0146 | 0.0000 | 6.0071 | 0.0000 | 0.0000 | 0.0493 |
| 184 | 4.3195 | 0.0699 | 1.0443 | 43.8566 | 0.0000 | 8.4918 | 0.0009 | 0.0162 | 0.0716 |
| 185 | 0.0000 | 0.0000 | 0.0000 | 0.0000 | 1.0000 | 0.0000 | 0.0000 | 0.0000 | 0.0000 |
| 186 | 0.1373 | 0.0000 | 0.0381 | 2.5665 | 0.1092 | 0.3095 | 0.0000 | 0.0000 | 0.7378 |
| 187 | 1.3529 | 0.0000 | 0.3201 | 18.1218 | 0.0000 | 2.6032 | 0.0000 | 0.0000 | 0.0888 |
| 188 | 0.0000 | 0.0000 | 0.0000 | 0.0000 | 1.0000 | 0.0000 | 0.0000 | 0.0000 | 0.0000 |
| 189 | 0.2720 | 0.0000 | 0.0441 | 3.6125 | 0.0573 | 0.3584 | 0.0000 | 0.0000 | 0.3735 |
| 190 | 1.8571 | 0.0000 | 0.3924 | 23.7914 | 0.0000 | 3.1908 | 0.0000 | 0.0000 | 0.0653 |
| 191 | 0.9044 | 0.0000 | 0.1654 | 9.9503 | 0.0016 | 1.3450 | 0.0000 | 0.0000 | 0.1449 |
| 192 | 0.0000 | 0.0000 | 0.0000 | 0.0000 | 1.0000 | 0.0000 | 0.0000 | 0.0000 | 0.0000 |
| 193 | 0.9156 | 0.0000 | 0.1660 | 10.0482 | 0.0015 | 1.3502 | 0.0000 | 0.0000 | 0.1431 |
| 194 | 0.2724 | 0.0000 | 0.0451 | 3.5786 | 0.0585 | 0.3665 | 0.0000 | 0.0000 | 0.3827 |
| 195 | 0.1373 | 0.0000 | 0.0392 | 2.5100 | 0.1131 | 0.3184 | 0.0000 | 0.0000 | 0.7677 |
| 196 | 0.8673 | 0.0531 | 0.1785 | 6.8492 | 0.0089 | 1.4516 | 0.0035 | 0.0612 | 0.2703 |
| 197 | 0.6237 | 0.0000 | 0.0904 | 7.7740 | 0.0053 | 0.7354 | 0.0000 | 0.0000 | 0.1619 |
| 198 | 0.7734 | 0.0000 | 0.1617 | 12.4902 | 0.0004 | 1.3145 | 0.0000 | 0.0000 | 0.1260 |
| 199 | 0.9136 | 0.0000 | 0.2416 | 15.6381 | 0.0001 | 1.9647 | 0.0000 | 0.0000 | 0.1160 |
| 200 | 0.0000 | 0.0439 | 0.0439 | 0.0088 | 0.9254 | 0.3566 | 1462.1862 | 10000.000<br>0 | 10000.000<br>0 |
| 201 | 0.8954 | 0.0000 | 0.2625 | 17.0957 | 0.0000 | 2.1349 | 0.0000 | 0.0000 | 0.1132 |
| 202 | 0.5743 | 0.0690 | 0.1664 | 3.1111 | 0.0778 | 1.3533 | 0.0068 | 0.1202 | 0.5326 |
| 203 | 3.0000 | 0.0000 | 0.8142 | 50.6222 | 0.0000 | 6.6207 | 0.0000 | 0.0000 | 0.0351 |
| 204 | 0.4000 | 0.0000 | 0.0825 | 6.2389 | 0.0125 | 0.6712 | 0.0000 | 0.0000 | 0.2512 |
| 205 | 1.2072 | 0.0537 | 0.2628 | 11.6100 | 0.0007 | 2.1373 | 0.0025 | 0.0445 | 0.1973 |
| 206 | 1.3529 | 0.0000 | 0.3132 | 21.2564 | 0.0000 | 2.5470 | 0.0000 | 0.0000 | 0.0750 |
| 207 | 0.2877 | 0.0000 | 0.0444 | 3.7112 | 0.0540 | 0.3610 | 0.0000 | 0.0000 | 0.3519 |
| 208 | 2.6056 | 0.0000 | 0.5748 | 39.9407 | 0.0000 | 4.6744 | 0.0000 | 0.0000 | 0.0376 |
| 209 | 0.5731 | 0.0000 | 0.1538 | 10.3900 | 0.0013 | 1.2509 | 0.0000 | 0.0000 | 0.1770 |
| 210 | 0.5502 | 0.0000 | 0.0759 | 7.9203 | 0.0049 | 0.6170 | 0.0000 | 0.0000 | 0.1531 |
| 211 | 0.0000 | 0.0000 | 0.0000 | 0.0000 | 1.0000 | 0.0000 | 0.0000 | 0.0000 | 0.0000 |
| 212 | 1.6439 | 0.0000 | 0.3321 | 24.7877 | 0.0000 | 2.7006 | 0.0000 | 0.0000 | 0.0609 |
| 213 | 0.3655 | 0.1001 | 0.1571 | 1.5333 | 0.2156 | 1.2773 | 0.0454 | 0.2740 | 0.8525 |
| 214 | 4.5596 | 0.0000 | 0.6112 | 53.3306 | 0.0000 | 4.9697 | 0.0000 | 0.0000 | 0.0213 |
| 215 | 0.0000 | 0.0000 | 0.0000 | 0.0000 | 1.0000 | 0.0000 | 0.0000 | 0.0000 | 0.0000 |
| 216 | 1.8153 | 0.0000 | 0.2159 | 20.6059 | 0.0000 | 1.7556 | 0.0000 | 0.0000 | 0.0522 |
| 217 | 0.9243 | 0.0000 | 0.2666 | 14.5720 | 0.0001 | 2.1675 | 0.0000 | 0.0000 | 0.1312 |
| 218 | 2.0769 | 0.0000 | 0.3184 | 21.3914 | 0.0000 | 2.5891 | 0.0000 | 0.0000 | 0.0590 |
| 219 | 0.6452 | 0.0000 | 0.0910 | 7.8870 | 0.0050 | 0.7397 | 0.0000 | 0.0000 | 0.1567 |
| 220 | 0.0928 | 0.0000 | 0.0395 | 1.1939 | 0.2746 | 0.3210 | 0.0000 | 0.0000 | 1.7623 |
| 221 | 0.0000 | 0.0000 | 0.0000 | 0.0000 | 1.0000 | 0.0000 | 0.0000 | 0.0000 | 0.0000 |
| 222 | 0.2835 | 0.0000 | 0.0789 | 5.0825 | 0.0242 | 0.6414 | 0.0000 | 0.0000 | 0.3719 |
| 223 | 2.3567 | 0.0000 | 0.3682 | 24.3183 | 0.0000 | 2.9942 | 0.0000 | 0.0000 | 0.0521 |
| 224 | 1.6066 | 0.0000 | 0.4605 | 22.0347 | 0.0000 | 3.7447 | 0.0000 | 0.0000 | 0.0858 |
| 225 | 11.2648 | 0.0000 | 1.0274 | 68.4271 | 0.0000 | 8.3540 | 0.0000 | 0.0000 | 0.0116 |
| 226 | 0.7404 | 0.0000 | 0.2181 | 12.0620 | 0.0005 | 1.7737 | 0.0000 | 0.0000 | 0.1610 |
| 227 | 0.1729 | 0.0000 | 0.0784 | 3.7735 | 0.0521 | 0.6379 | 0.0000 | 0.0000 | 0.7082 |
| 228 | 1.7586 | 0.0000 | 0.4540 | 30.8744 | 0.0000 | 3.6920 | 0.0000 | 0.0000 | 0.0593 |
| 229 | 2.3195 | 0.0000 | 0.6722 | 35.6279 | 0.0000 | 5.4661 | 0.0000 | 0.0000 | 0.0530 |
| 230 | 0.4314 | 0.0000 | 0.1189 | 7.6758 | 0.0056 | 0.9671 | 0.0000 | 0.0000 | 0.2444 |
| 231 | 0.0000 | 0.0000 | 0.0000 | 0.0000 | 1.0000 | 0.0000 | 0.0000 | 0.0000 | 0.0000 |
| 232 | 0.0000 | 0.0000 | 0.0000 | 0.0000 | 1.0000 | 0.0000 | 0.0000 | 0.0000 | 0.0000 |
| 233 | 0.5711 | 0.1372 | 0.1973 | 2.0310 | 0.1541 | 1.6047 | 0.0597 | 0.2403 | 0.6289 |
| 234 | 0.0000 | 0.0000 | 0.0000 | 0.0000 | 1.0000 | 0.0000 | 0.0000 | 0.0000 | 0.0000 |
| 235 | 0.2636 | 0.0000 | 0.0440 | 3.5732 | 0.0587 | 0.3576 | 0.0000 | 0.0000 | 0.3853 |
| 236 | 2.6511 | 0.0000 | 0.7753 | 46.0604 | 0.0000 | 6.3044 | 0.0000 | 0.0000 | 0.0390 |
| 237 | 1.0663 | 0.0560 | 0.3290 | 12.7008 | 0.0004 | 2.6757 | 0.0030 | 0.0525 | 0.2331 |
| 238 | 0.6357 | 0.0000 | 0.1581 | 11.0293 | 0.0009 | 1.2852 | 0.0000 | 0.0000 | 0.1594 |
| 239 | 0.0000 | 0.0000 | 0.0000 | 0.0000 | 1.0000 | 0.0000 | 0.0000 | 0.0000 | 0.0000 |
| 240 | 2.0993 | 0.0000 | 0.8145 | 34.8044 | 0.0000 | 6.6232 | 0.0000 | 0.0000 | 0.0635 |
| 241 | 0.6014 | 0.0000 | 0.0939 | 7.3236 | 0.0068 | 0.7636 | 0.0000 | 0.0000 | 0.1788 |
| 242 | 1.3529 | 0.0000 | 0.1970 | 20.0704 | 0.0000 | 1.6019 | 0.0000 | 0.0000 | 0.0637 |
| 243 | 0.4994 | 0.0000 | 0.1160 | 8.6482 | 0.0033 | 0.9429 | 0.0000 | 0.0000 | 0.1943 |
| 244 | 0.0000 | 0.0000 | 0.0000 | 0.0000 | 1.0000 | 0.0000 | 0.0000 | 0.0000 | 0.0000 |
| 245 | 0.0000 | 0.0000 | 0.0000 | 0.0000 | 1.0000 | 0.0000 | 0.0000 | 0.0000 | 0.0000 |
| 246 | 0.0928 | 0.0000 | 0.0366 | 2.1737 | 0.1404 | 0.2972 | 0.0000 | 0.0000 | 1.1363 |
| 247 | 0.7803 | 0.0000 | 0.2206 | 12.6274 | 0.0004 | 1.7937 | 0.0000 | 0.0000 | 0.1503 |
| 248 | 0.1649 | 0.0000 | 0.0395 | 2.8905 | 0.0891 | 0.3211 | 0.0000 | 0.0000 | 0.6024 |
| 249 | 1.0218 | 0.1248 | 0.4662 | 10.2373 | 0.0014 | 3.7913 | 0.0202 | 0.1222 | 0.3798 |
| 250 | 0.0000 | 0.0000 | 0.0000 | 0.0000 | 1.0000 | 0.0000 | 0.0000 | 0.0000 | 0.0000 |
| 251 | 0.0000 | 0.0000 | 0.0000 | 0.0000 | 1.0000 | 0.0000 | 0.0000 | 0.0000 | 0.0000 |
| 252 | 0.0000 | 0.0000 | 0.0000 | 0.0000 | 1.0000 | 0.0000 | 0.0000 | 0.0000 | 0.0000 |
| 253 | 0.1668 | 0.0000 | 0.0394 | 2.8836 | 0.0895 | 0.3208 | 0.0000 | 0.0000 | 0.5956 |
| 254 | 0.2793 | 0.1183 | 0.1771 | 0.8135 | 0.3671 | 1.4401 | 0.0704 | 0.4234 | 1.3076 |
| 255 | 1.2093 | 0.0823 | 0.5111 | 11.5180 | 0.0007 | 4.1560 | 0.0039 | 0.0680 | 0.3022 |
| 256 | 0.1388 | 0.0000 | 0.0395 | 2.5133 | 0.1129 | 0.3213 | 0.0000 | 0.0000 | 0.7658 |
| 257 | 0.0000 | 0.0000 | 0.0000 | 0.0000 | 1.0000 | 0.0000 | 0.0000 | 0.0000 | 0.0000 |
| 258 | 1.9630 | 0.0000 | 0.4167 | 30.3846 | 0.0000 | 3.3887 | 0.0000 | 0.0000 | 0.0513 |
| 259 | 0.9679 | 0.0000 | 0.2005 | 15.4029 | 0.0001 | 1.6302 | 0.0000 | 0.0000 | 0.1003 |
| 260 | 0.2929 | 0.0000 | 0.1083 | 5.9411 | 0.0148 | 0.8807 | 0.0000 | 0.0000 | 0.3766 |
| 261 | 0.1136 | 0.0508 | 0.0703 | 0.3166 | 0.5737 | 0.5720 | 0.0255 | 0.4468 | 1.9804 |

These results are also available at the following link: https://github.com/aglucaci/AnalysisOfOrthologousCollections/blob/main/tables/BDNF/BDNF_FEL_CI.csv

## REFERENCES

1. Notaras, M., R. Hill, and M. van den Buuse, The BDNF gene Val66Met polymorphism as a modifier of psychiatric disorder susceptibility: progress and controversy. Mol Psychiatry, 2015. 20(8): p. 916–930.

2. Nagappan, G. and B. Lu, Activity-dependent modulation of the BDNF receptor TrkB: mechanisms and implications. Trends in Neurosci, 2005. 28(9): p. 464–471.

3. Black, I.B., Trophic regulation of synaptic plasticity. J Neurobiol, 1999. 41(1): p. 108–118.

4. Yoshii, A. and M. Constantine-Paton, Postsynaptic BDNF-TrkB signaling in synapse maturation, plasticity, and disease. Dev Neurobiol, 2010. 70(5): p. 304–322.

5. Sakuragi, S., K. Tominaga-Yoshino, and A. Ogura, Involvement of TrkB-and p75 NTR-signaling pathways in two contrasting forms of long-lasting synaptic plasticity. Sci Reps, 2013. 3(1): p. 1–7.

6. Horch, H.W., et al., Destabilization of cortical dendrites and spines by BDNF. Neuron, 1999. 23(2): p. 353–364.

7. Giza, J.I., et al., The BDNF Val66Met prodomain disassembles dendritic spines altering fear extinction circuitry and behavior. Neuron, 2018. 99(1): p. 163–178. E6.

8. Martinowich, K., H. Manji, and B. Lu, New insights into BDNF function in depression and anxiety. Nature Neurosci, 2007. 10(9): p. 1089–1093.

9. Angelucci, F., S. Brene, and A. Mathe, BDNF in schizophrenia, depression and corresponding animal models. Mol Psychiatry, 2005. 10(4): p. 345–352.

10. Kim, Y.-K., et al., Low plasma BDNF is associated with suicidal behavior in major depression. Prog Neuropsychopharmacol Biol Psychiatry, 2007. 31(1): p. 78–85.

11. Notaras, M. and M. van den Buuse, Neurobiology of BDNF in fear memory, sensitivity to stress, and stress-related disorders. Mol Psychiatry, 2020. 25(10): p. 2251–2274.

12. Pivac, N., et al., The association between brain-derived neurotrophic factor Val66Met variants and psychotic symptoms in posttraumatic stress disorder. World J Biol Psychiatry, 2012. 13(4): p. 306–311.

13. Pitts, B.L., et al., BDNF Val66Met polymorphism and posttraumatic stress symptoms in US military veterans: Protective effect of physical exercise. Psychoneuroendocrinol, 2019. 100: p. 198–202.

14. Zhang, L., et al., PTSD risk is associated with BDNF Val66Met and BDNF overexpression. Mol Psychiatry, 2014. 19(1): p. 8–10.

15. Notaras, M., R. Hill, and M. Van den Buuse, A role for the BDNF gene Val66Met polymorphism in schizophrenia? A comprehensive review. Neurosci Biobehav Rev, 2015. 51: p. 15–30.

16. Gratacòs, M., et al., Brain-derived neurotrophic factor Val66Met and psychiatric disorders: meta-analysis of case-control studies confirm association to substance-related disorders, eating disorders, and schizophrenia. Biol Psychiatry, 2007. 61(7): p. 911–922.

17. Zakharyan, R., et al., Functional variants of the genes involved in neurodevelopment and susceptibility to schizophrenia in an Armenian population. Human Immunol, 2011. 72(9): p. 746–748.

18. Howells, D., et al., Reduced BDNF mRNA expression in the Parkinson’s disease substantia nigra. Exp Neurol, 2000. 166(1): p. 127–135.

19. Palasz, E., et al., BDNF as a promising therapeutic agent in Parkinson’s disease. Int J Mol Sci, 2020. 21(3): p. 1170.

20. Correia, C., et al., Increased BDNF levels and NTRK2 gene association suggest a disruption of BDNF/TrkB signaling in autism. Genes, Brain, Behav, 2010. 9(7): p. 841–848.

21. Ricci, S., et al., Altered cytokine and BDNF levels in autism spectrum disorder. Neurotox Res, 2013. 24(4): p. 491–501.

22. Tsai, S.-J., Is autism caused by early hyperactivity of brain-derived neurotrophic factor? Med Hypotheses, 2005. 65(1): p. 79–82.

23. Massa, S.M., et al., Small molecule BDNF mimetics activate TrkB signaling and prevent neuronal degeneration in rodents. J Clin Investig, 2010. 120(5): p. 1774–1785.

24. Kingwell, K., BDNF copycats. Nature Rev Drug Discov, 2010. 9(6): p. 433–433

25. Chen, B., et al., Increased hippocampal BDNF immunoreactivity in subjects treated with antidepressant medication. Biol Psychiatry, 2001. 50(4): p. 260–265.

26. Björkholm, C. and L.M. Monteggia, BDNF–a key transducer of antidepressant effects. Neuropharmacol, 2016. 102: p. 72–79.

27. Anastasia, A., et al., Val66Met polymorphism of BDNF alters prodomain structure to induce neuronal growth cone retraction. Nat Comm, 2013. 4(1): p. 1–13.

28. Glerup, S., et al., SorCS2 is required for BDNF-dependent plasticity in the hippocampus. Mol Psychiatry, 2016. 21(12): p. 1740–1751.

29. Notaras, M. and M. van den Buuse, Brain-derived neurotrophic factor (BDNF): novel insights into regulation and genetic variation. The Neuroscientist, 2019. 25(5): p. 434–454.

30. Pruunsild, P., et al., Dissecting the human BDNF locus: bidirectional transcription, complex splicing, and multiple promoters. Genomics, 2007. 90(3): p. 397–406.

31. Lipovich, L., et al., Activity-dependent human brain coding/noncoding gene regulatory networks. Genetics, 2012. 192(3): p. 1133–1148.

32. Chiaruttini, C., et al., Dendritic trafficking of BDNF mRNA is mediated by translin and blocked by the G196A (Val66Met) mutation. Proc Nat Acad Sci USA, 2009. 106(38): p. 16481–16486.

33. Chen, Z.-Y., et al., Sortilin controls intracellular sorting of brain-derived neurotrophic factor to the regulated secretory pathway. J Neurosci, 2005. 25(26): p. 6156–6166.

34. Gray, K. and V. Ellis, Activation of pro-BDNF by the pericellular serine protease plasmin. FEBS letters, 2008. 582(6): p. 907–910.

35. del Carmen Cardenas-Aguayo, M., et al., Neurogenic and neurotrophic effects of BDNF peptides in mouse hippocampal primary neuronal cell cultures. PloS One, 2013. 8(1): p. E53596.

36. Maisonpierre, P.C., et al., Human and rat brain-derived neurotrophic factor and neurotrophin-3: gene structures, distributions, and chromosomal localizations. Genomics, 1991. 10(3): p. 558–568.

37. Finn, R.D., et al., The Pfam protein families database: towards a more sustainable future. Nucleic Acids Research, 2016. 44(D1): p. D279–D285.

38. Rodriguez-Tebar, A., G. Dechant, and Y.-A. Barde, Binding of brain-derived neurotrophic factor to the nerve growth factor receptor. Neuron, 1990. 4(4): p. 487–492.

39. Castellani, V. and J. Bolz, Opposing roles for neurotrophin-3 in targeting and collateral formation of distinct sets of developing cortical neurons. Development, 1999. 126(15): p. 3335–3345.

40. Kuczewski, N., et al., Activity-dependent dendritic release of BDNF and biological consequences. Mol Neurobiol, 2009. 39(1): p. 37–49.

41. Chen, Z.-Y., et al., Variant brain-derived neurotrophic factor (BDNF)(Met66) alters the intracellular trafficking and activity-dependent secretion of wild-type BDNF in neurosecretory cells and cortical neurons. J Neurosci, 2004. 24(18): p. 4401–4411.

42. Kumar, S., et al., TimeTree: a resource for timelines, timetrees, and divergence times. Mol Biol Evol, 2017. 34(7): p. 1812–1819.

43. Hedges, S.B., J. Dudley, and S. Kumar, TimeTree: a public knowledge-base of divergence times among organisms. Bioinformatics, 2006. 22(23): p. 2971–2972.IQ-TREE reference.

44. Sergei L Kosakovsky Pond, Art FY Poon, Ryan Velazquez, Steven Weaver, N Lance Hepler, Ben Murrell, Stephen D Shank, Brittany Rife Magalis, Dave Bouvier, Anton Nekrutenko, Sadie Wisotsky, Stephanie J Spielman, Simon DW Frost, Spencer V Muse (2020) HyPhy 2.5—A Customizable Platform for Evolutionary Hypothesis Testing Using Phylogenies. 1 Molecular Biology and Evolution 37.1 (2020): 295-299.

45. Martin DP, Murrell B, Golden M, Khoosal A, & Muhire B (2015) RDP4: Detection and analysis of recombination patterns in virus genomes. Virus Evolution 1: vev003 doi: 10.1093/ve/vev003.

46. B.Q. Minh, H.A. Schmidt, O. Chernomor, D. Schrempf, M.D. Woodhams, A. von Haeseler, R. Lanfear (2020) IQ-TREE 2: New models and efficient methods for phylogenetic inference in the genomic era. Mol. Biol. Evol., 37:1530–1534. https://doi.org/10.1093/molbev/msaa015

47. Sergei L. Kosakovsky Pond and Simon D. W. Frost (2005) Not So Different After All: A Comparison of Methods for Detecting Amino Acid Sites Under Selection Molecular Biology and Evolution 22(5): 1208–1222

48. Wisotsky SR, Kosakovsky Pond SL, Shank SD, Muse SV. Synonymous Site-to-Site Substitution Rate Variation Dramatically Inflates False Positive Rates of Selection Analyses: Ignore at Your Own Peril. Mol Biol Evol. 2020 Aug 1;37(8):2430–2439. doi: 10.1093/molbev/msaa037. PMID: 32068869; PMCID: PMC7403620.

49. Ben Murrell, Joel O. Wertheim, Sasha Moola, Thomas Weighill, Konrad Scheffler and Sergei L. Kosakovsky Pond (2012) Detecting Individual Sites Subject to Episodic Diversifying Selection PLoS Genetics 8(7): e1002764.

50. M. D. Smith, J. O. Wertheim, S. Weaver, B. Murrell, K. Scheffler and S. L. Kosakovsky Pond Less Is More: An Adaptive Branch-Site Random Effects Model for Efficient Detection of Episodic Diversifying Selection Molecular Biology and Evolution 32: 1342–1353.

51. Art F. Y. Poon, Fraser I. Lewis, Simon D. W. Frost, Sergei L. Kosakovsky Pond, Spidermonkey: rapid detection of co-evolving sites using Bayesian graphical models, Bioinformatics, Volume 24, Issue 17, 1 September 2008, Pages 1949–1950, https://doi.org/10.1093/bioinformatics/btn313.

52. Wertheim JO, Murrell B, Smith MD, Kosakovsky Pond SL, Scheffler K. RELAX: detecting relaxed selection in a phylogenetic framework. Mol Biol Evol. 2015 Mar;32(3):820–32. doi: 10.1093/molbev/msu400. Epub 2014 Dec 23. PMID: 25540451; PMCID: PMC4327161.

53. Kosakovsky Pond SL, Wisotsky SR, Escalante A, Magalis BR, Weaver S. Contrast-FEL- A Test for Differences in Selective Pressures at Individual Sites among Clades and Sets of Branches. Mol Biol Evol. 2021 Mar 9;38(3):1184–1198. doi: 10.1093/molbev/msaa263. PMID: 33064823; PMCID: PMC7947784.

54. Lucaci, Alexander G., Sadie R. Wisotsky, Stephen D. Shank, Steven Weaver, and Sergei L. Kosakovsky Pond. “Extra Base Hits: Widespread Empirical Support for Instantaneous Multiple-Nucleotide Changes.” PLOS ONE 16, no. 3 (March 12, 2021): e0248337. https://doi.org/10.1371/journal.pone.0248337.

55. Martin, D. & Rybicki, E. (2000). RDP: detection of recombination amongst aligned sequences. Bioinformatics 16, 562–563.

56. Padidam, M., Sawyer, S. & Fauquet, C. M. (1999). Possible emergence of new geminiviruses by frequent recombination. Virology 265, 218–225.

57. Posada, D. & Crandall, K. A. (2001). Evaluation of methods for detecting recombination from DNA sequences: Computer simulations. Proc Natl Acad Sci 98, 13757–13762.

58. Maynard Smith, J. (1992). Analyzing the mosaic structure of genes. J Mol Evol 34, 126–129.

59. Martin, D. P., Posada, D., Crandall, K. A. & Williamson, C. (2005). A modified bootscan algorithm for automated identification of recombinant sequences and recombination breakpoints. AIDS Res Hum Retroviruses 21, 98–102.

60. Gibbs, M. J., Armstrong, J. S. & Gibbs, A. J. (2000). Sister-Scanning: a Monte Carlo procedure for assessing signals in recombinant sequences. Bioinformatics 16, 573–582.

61. Lam H.M., Ratmann O., Boni M.F. (2018). Improved algorithmic complexity for the 3SEQ recombination detection algorithm. Mol Biol Evol, 35, 247–251.

